# Brainstem enkephalinergic neural circuit underlying cold-induced pain relief in mice

**DOI:** 10.1101/2025.11.10.686886

**Authors:** Hayun kim, Yoonkyung Lee, Seog Bae oh

**Author notes:** Corresponding Author: Seog Bae Oh, DDS, PhD.

## Abstract

Application of cold or cold-mimicking chemicals to injury has long been recognized as an effective means of pain relief and is widely utilized in daily life. However, underlying neural mechanisms remain elusive. Here, we identified a cold-responsive neuronal subset within lateral parabrachial nucleus (lPBN), the thermosensory relay region in the hindbrain, that mediates cold-induced analgesia in mice. Selective activation of these neurons and their projection to ventrolateral periaqueductal gray (vlPAG) increased nociceptive threshold via opioid receptor signaling in vlPAG. Conversely, ablation of these neurons attenuated analgesia induced by cold-mimicking chemicals. We further identified that these neurons express precursor gene of enkephalin, which is released into vlPAG for pain relief. Activation of cold-responsive neurons in descending pain modulation circuitry reduced spinal cord responses to noxious stimuli, suggesting the involvement of top-down pain modulation pathway. These findings propose a central mechanism underlying cold-induced pain relief, which could be a novel therapeutic target.

**Teaser:** Cold-sensitive lPBN neurons release enkephalin into vlPAG for top-down analgesic action during cold-induced pain relief

## Introduction

Cooling of the regions associated with pain has long been recognized as a method to induce pain relief (analgesia) (*1–6*). Historically, Hippocrates documented the use of cold water to alleviate joint pain, ulcers, and sprains approximately 2,400 years ago (*7*). Similarly, administration of a chemical agonist for transient receptor potential melastatin 8 (TRPM8), which is an essential ion channel involved in cooling perception (*8, 9*), induces robust analgesia, mimicking cold-induced activation of TRPM8 channel in peripheral sensory neurons. In contemporary clinical practice, over-the-counter creams and transdermal patches containing the TRPM8 agonists such as menthol or eucalyptol are widely utilized as the medication for pain management (*2, 3*).

Such cold-induced, TRPM8-mediated analgesia, has been attributed to peripheral mechanism, including decreased peripheral nerve conductance by physical cooling, desensitization of nociceptive nerve endings and inactivation of voltage gated calcium and/or sodium channels by menthol (*6, 10, 11*). Nevertheless, multiple lines of evidence suggest that cold-induced analgesia is primarily mediated by the central nervous system rather than by peripheral mechanisms. For instance, genetic ablation of cold-sensitive TRPM8^+^ fibers, but not nociceptor, abolished cold-induced analgesia (*12*). This results suggest that cold-induced analgesia does not originate from a direct inhibition of nociceptor activity upon cold application, but rather involves an inhibition of nociceptive signaling by TRPM8^+^ fibers at the central level. Additionally, pain relief induced by skin cooling and menthol is significantly attenuated by the opioid receptor antagonist naloxone both in human subjects and preclinical mouse models, indicating the involvement of a central endogenous opioidergic pain modulation mechanism in cooling-induced analgesia (*1, 4, 13*). Moreover, cold-induced analgesia is not confined to the side of cold application; stimulation of a distal or even contralateral regions similarly elicited marked analgesic effects (*14, 15*).

Temperature information from periphery is detected via thermosensory neurons and subsequently transmitted to superficial dorsal horn of spinal cord and supraspinal regions for temperature information processing and representation (*16*). Among the supraspinal regions involved in temperature representation, the lateral parabrachial nucleus (lPBN) is well recognized as a first-order sensory gateway, integrating nociceptive and thermosensory signals including cold (*17, 18*). Through various interoceptive and external signaling, lPBN also functions as a regulatory node for pain based on homeostatic demands (*19–21*). Indeed, recent studies have highlighted the crucial role of the lPBN in pain regulation. For instance, activating specific subset of lPBN neurons induces acute analgesia and reversed morphine-induced hyperalgesia (*22, 23*). Especially, activation of the lPBN to periaqueductal gray (PAG) neural circuit has been implicated in analgesic processes (*24, 25*). Thus, it is plausible that the lPBN and its downstream projections might contribute to cold-induced analgesia as a top-down pain modulation mechanism. Collectively, these findings suggest that peripheral cold stimulation elicits analgesic effects, which are dominantly mediated by central nervous system pathways, extending beyond local or peripheral effects. However, despite the historical and clinical use of cold-induced analgesia, the central neural circuits and molecular mechanism underlying such analgesic effect has been insufficiently understood.

To address this gap in knowledge, we employed integrative approaches combining behavioral assays in mouse model, chemogenetic activity modulation using permanent marking of active neural clusters through targeted recombination in active neural populations (FosTRAP), *in vivo* calcium imaging and transcriptomic analysis to elucidate the circuit-to-molecular mechanism of cold-induced analgesia. Notably, we identified a population of cold-responsive neurons within the lPBN (lPBN^cold^ neurons). lPBN^cold^ neurons were found to be necessary and sufficient for TRPM8-dependent analgesia, mediating analgesic effect via projections to the ventrolateral PAG (vlPAG). Furthermore, activation of lPBN^cold^ neurons released enkephalin, a well-known antinociceptive neuropeptide, and diminishes noxious stimulation-induced Fos in the spinal cord, suggesting engagement of the descending inhibitory pathways. These findings provide novel insights into the neural circuits underlying cold-induced analgesia and highlight a critical role of lPBN-to-vlPAG neural circuits in TRPM8-depenedent analgesia.

## Results

### TRPM8-mediated antinociception and activation of supraspinal brain regions by cooling

To investigate whether cutaneous cooling elicits pain relief (analgesia) in mice model, we first assessed the effects of peripheral cold stimulation on mechanical pain sensitivity by exposing hindpaw to 17°C metal plate for 5 minutes (**fig. S1A**). As expected, physically cooling the hindpaw induced significant analgesia in Complete Freund’s adjuvant (CFA)-induced inflammatory pain model, in both mechanical pain and heat pain (**fig. S1B, C**). Furthermore, to avoid the change of peripheral sensory nerve conductance under physical cooling and given the technical limitations of simultaneously applying physical cooling and conducting behavioral assays in the same hindpaw for a prolonged time period (*26*), we utilized topical application of potent TRPM8 agonist icilin (**Fig. 1A**), which mimic cooling of skin through activation of TRPM8-expressing fiber as a behavioral testing paradigm (*27*). Consistent with the previous report (*28*), topical icilin treatment to hindpaw significantly increased mechanical withdrawal threshold in both naïve and CFA-induced inflammatory pain model (**Fig. 1B, C**). Furthermore, in concordance with the reports in human subjects (*3, 14, 15*), icilin treatment to distal regions (e.g., to the tail) resulted in an increased mechanical withdrawal threshold in the inflamed hindpaw (**Fig. 1D**). Since TRPM8-induced analgesia has traditionally been attributed to peripheral mechanism such as desensitization or inhibition of voltage gated calcium channels (*6, 10*), we first aimed to investigate the effect of TRPM8 activation within the peripheral nervous system. To assess this, we repeatedly applied KCl to primary-cultured dorsal root ganglion neurons from mice to induce membrane depolarization challenge and measure the magnitude of calcium response (*29*). We then tested whether KCl-mediated calcium response could be altered by icilin treatment (**fig. S2**). Our analysis revealed no significant difference in the desensitization ratio between icilin-responsive (presumed TRPM8^+^) and icilin-nonresponsive neurons (presumed TRPM8^-^). These findings suggest a lack of neuronal desensitization through TRPM8 activation, highlighting the involvement of central nervous system, but not peripheral nervous system in cold-induced analgesia.

**Figure 1.**
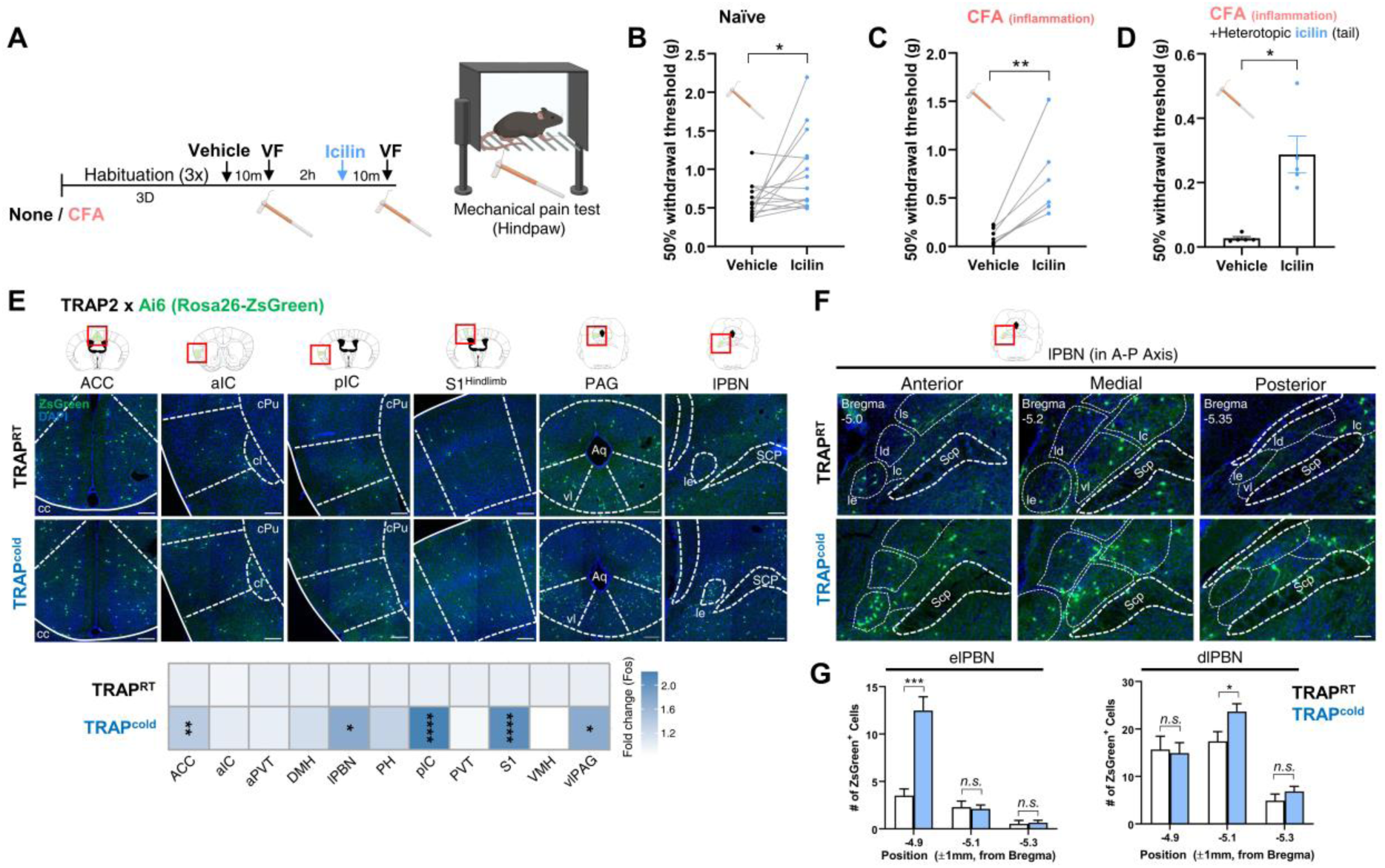
TRPM8-mediated antinociception and activation of supraspinal brain regions by cooling. (A) Experimental scheme and timeline for the icilin-induced pain relief experiment. (B) Mechanical sensitivity measure with 50% withdrawal thresholds in naïve mice in topical vehicle- or icilin-treatment. (C) Ipsilateral paw mechanical sensitivity measure with 50% withdrawal thresholds in CFA-induced inflammatory pain model mice in topical vehicle- or icilin-treatment. (D) Mechanical sensitivity measure with 50% withdrawal thresholds in naïve mice in topical vehicle- or icilin-treatment to tail (dstal) region. (E) Distribution of Zsg-positive neurons in a diverse brain region of a control-TRAP mouse and Cold-TRAP mouse (top) and quantification of Zsg-positive neurons neurons per section (normalized to control; bottom). Green: ZsGreen, Blue: DAPI. Scale bar: 100μm. (F) Distribution of Zsg-positive neurons of a control-TRAP mouse and Cold-TRAP mouse in lateral parabrachial nucleus. Green: ZsGreen, Blue: DAPI. Scale bar: 100μm. (G) Quantification of Zsg-positive neurons in external lateral division of lateral parabrachial nucleus (elPBN, left) and dorsolateral division of lateral parabrachial nucleus (dlPBN, right). Data are presented as mean ± SEMs. ∗*p* < 0.05, ∗∗*p* < 0.01, ∗∗∗*p* < 0.001 and ∗∗∗∗*p* < 0.0001. Number of mice and statistical tests are listed in **Table S1**.

Considering prior evidence implicating supraspinal structures in cold-induced analgesia (*14, 30*), we next sought to characterize the pattern of neuronal activation at the whole-brain level. To selectively label and further manipulate neurons responsive to peripheral cold stimulus, we utilized FosTRAP transgenic mice (Fos-iCre/ERT2) crossed with Ai6 reporter mice (TRAP2; Ai6). Consistent with previous reports (*31, 32*), we observed robust peripheral cold stimulus-induced neuronal activation in the posterior insular cortex (pIC) and primary somatosensory cortex (S1), regions known to be activated by non-noxious cooling and TRPM8 agonist exposure. Notably, we also detected significant increase in the number of labeled neurons within the lPBN (**Fig. 1E**), which has been suggested as primary sensory hub for peripheral sensory stimuli (*33*). Thus, we focused on the functional involvement of lPBN in the cold-induced analgesia in the following experiments. Given the anatomical and functional heterogeneity of the lPBN, which consists of multiple subnuclei with distinctive molecular and functional profiles (*34*), we quantified cold-responsive neurons along the rostrocaudal axis and within the specific subdivisions of the lPBN. Notably, Cold-responsive lPBN neurons were mainly located in rostral external lateral vision of lPBN (elPBN) and medial dorsal lateral division of lPBN (dlPBN) (**Fig. 1F, G**), consistent with a prior report (*17*). To note, cold-responsive lPBN neurons showed little overlap with calcitonin gene-related peptide (CGRP) through rostrocaudal axis (**fig. S3**), which constitutes a major population in medial-to-caudal elPBN (*34*).

### Response characteristics of lateral parabrachial nucleus neurons through *in vivo* calcium imaging

We utilized *in vivo* microendoscope to measure neuronal calcium response in the lPBN to investigate the real-time response patterns of lPBN to the array of peripheral stimulus *in vivo*. The activity of lPBN neurons at a single-cell level were monitored during peripheral stimulation using heat (55°C hot water), noxious pinch, and cold (4°C cold water) applied to the tail region (**Fig. 2A-C**), considering the lack of somatotopic arrangement in lPBN to peripheral stimulus (*35, 36*). *In vivo* calcium imaging revealed distinct neuronal populations within the lPBN: One subset exhibited responsiveness to heat and noxious pinch, but not responsive to cold stimuli (hereafter referred to as lPBN^HP^ (Heat-Pinch) neurons), while another subset is responsive to cold stimuli being (lPBN^cold^ neurons) but not to heat and noxious pinch stimuli. lPBN^HP^ and lPBN^cold^ neurons comprised 38.4% and 26.9% of total recorded neurons, respectively (**Fig. 2D**). Significant calcium response of lPBN^HP^ neurons, but not of lPBN^cold^ neurons, were observed 3.08 ± 0.19 seconds after the onset of prolonged heat exposure, corresponding to the average latency of the nocifensive tail-flick response (**Fig. 2E**). Similarly, tail pinch with blunt forceps induced robust calcium responses in lPBN^HP^ neurons, but not in lPBN^cold^ neurons (**Fig. 2F**). Conversely, cold stimulation significantly increased calcium response in lPBN^cold^ neurons while concomitantly decreasing calcium response of lPBN^HP^ neurons (**Fig. 2G**). To validate the functional segregation of lPBN neurons, we conducted double-labeling with TRAP/Fos immunostaining (**fig. S4**). Quantification of the TRAP method demonstrated a specificity of 0.50 and an efficiency of 0.35, further supporting the functional distinction between heat- and cold-responsive lPBN neuronal populations (**fig. S4B,** See **Table S1**). Cold-TRAP neurons were primarily localized to the rostral elPBN and medial dlPBN, whereas hot-TRAP neurons were predominantly found in the caudal elPBN and medial dlPBN. However, distribution of cold- and hot-responsive dlPBN neurons were not overlapping (**fig. S4C-E;** See **Table S1**).

**Figure 2.**
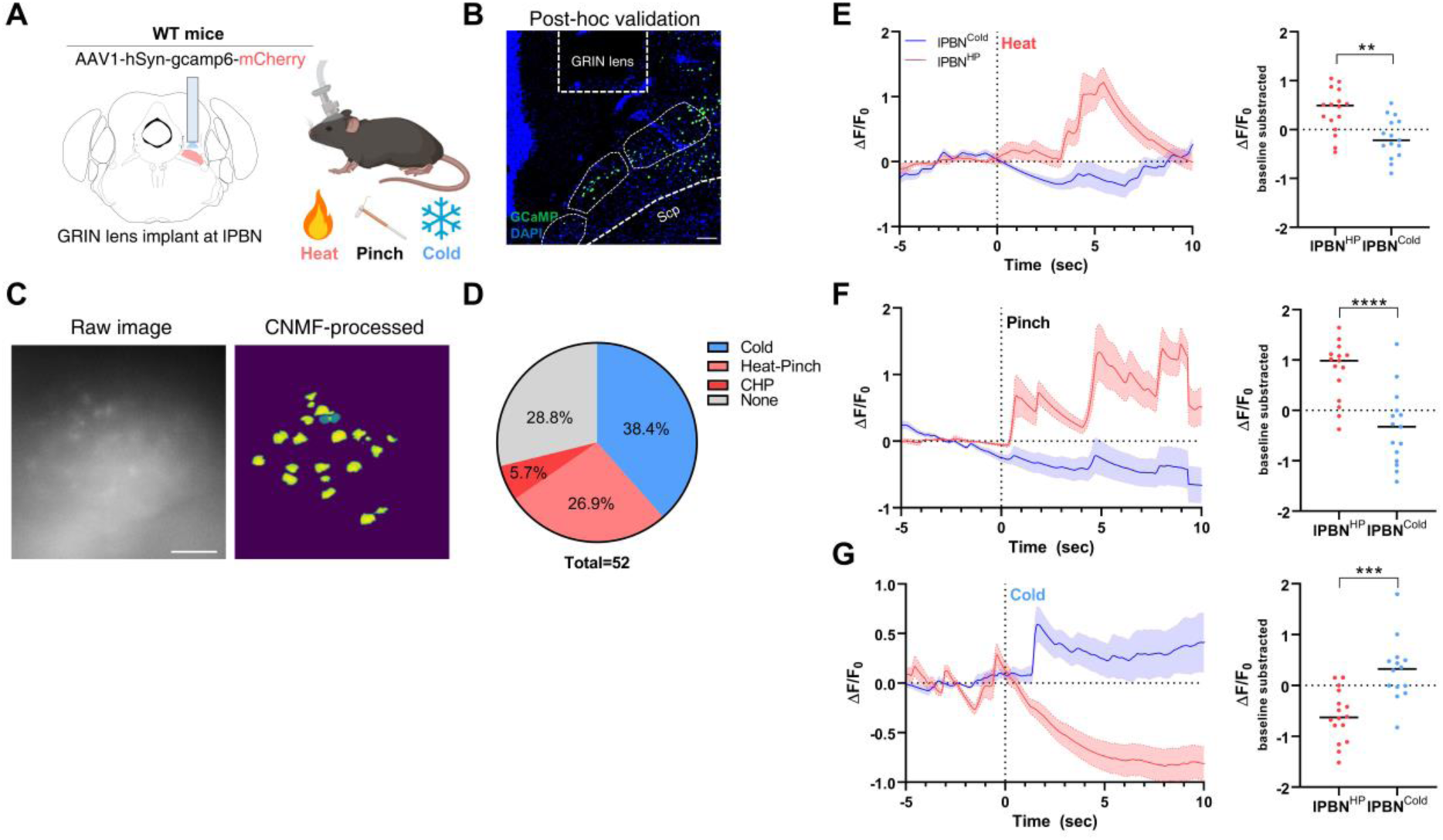
Response characteristics of lateral parabrachial nucleus neurons through *in vivo* calcium imaging. (A) Experimental scheme for the *in vivo* calcium imaging in lateral parabrachial nucleus. (B) Examples of virus expression and location of GRIN lens implantation. Green: GCaMP6f, Blue: DAPI. Scale bar: 100μm. (C) Examples of single cell imaging window in lateral parabrachial nucleus, both raw image from miniscope recording (left) and CNMF-processes image (right). Scale bar: 100μm. (D) Pie chart for cell population responsiveness in lPBN. (E) Peri-event plot of fluorescent signal recorded in lPBN neurons in response to heat stimuli. Responses of neurons are averaged by group and represented as mean ± SEMs (left) and quantification of average fluorescence difference from baseline (right). (F) Peri-event plot of fluorescent signal recorded in lPBN neurons in response to pinch stimuli. Responses of neurons are averaged by group and represented as mean ± SEMs (left) and quantification of average fluorescence difference from baseline (right). (G) Peri-event plot of fluorescent signal recorded in lPBN neurons in response to cold stimuli. Responses of neurons are averaged by group and represented as mean ± SEMs (left) and quantification of average fluorescence difference from baseline (right). Data are presented as mean ± SEMs. ∗*p* < 0.05, ∗∗*p* < 0.01, ∗∗∗*p* < 0.001 and ∗∗∗∗*p* < 0.0001. Number of mice and statistical tests are listed in **Table S1**.

### Chemogenetic activation of lPBN^cold^ neuron induces acute antinociception

To test the causal role of lPBN^cold^ neurons in analgesia, we first selectively activated lPBN^cold^ neurons using TRAP2 transgenic mice with excitatory DREADD hM3Dq (**Fig. 3A**). Using electrophysiological recordings in acute brain slices, we confirmed that treatment with Clozapine-N-oxide (CNO) significantly increased the activity of lPBN^cold^ neurons (**Fig. 3B–C, fig S5**). Notably, we observed that 70.5% of recorded lPBN^cold^ neurons exhibited spontaneous activity (**Fig. 3D**), an observation that contrasts with previous reports suggesting that only a small subset of lateral parabrachial nucleus (lPBN) neurons exhibit spontaneous action potential firing (20∼22% in both mouse and rat) (*37–39*). CNO-induced activation of lPBN^cold^ neurons elicited both tonic (11/12 cells) and burst (1/12 cells) firing patterns (**fig. S5D**). However, the resting membrane potentials of quiescent cells and spontaneously active cells did not significantly differ (**Fig. 3E**). Activation of lPBN^cold^ neurons with CNO injection significantly increased mechanical withdrawal threshold (**Fig. 3F**) and prolonged response latency to noxious hot (**Fig. 3G**) in a dose-dependent manner, while decreasing response latency to noxious cold (**Fig. 3H, I**). Expression level of DREAAD to bilateral lPBN correlates with CNO-induced antinociception, further supporting the role of lPBN neurons in antinociception (**Fig. 3J**). To further assess the behavioral effects of lPBN^cold^ neuron activation, we evaluated spontaneous exploration of thermal gradient plate. Activation of lPBN^cold^ neuron significantly reduced spontaneous entry into both noxious cold (below 17°C) and innocuous cold (between 17°C and 24°C) zones in thermal gradient plates, indicating heightened cold sensitivity during lPBN^cold^ neurons activation (**fig. S6B-E**). Nonetheless, activation of lPBN^cold^ neurons does not alter core body temperature (**fig. S6F**).

**Figure 3.**
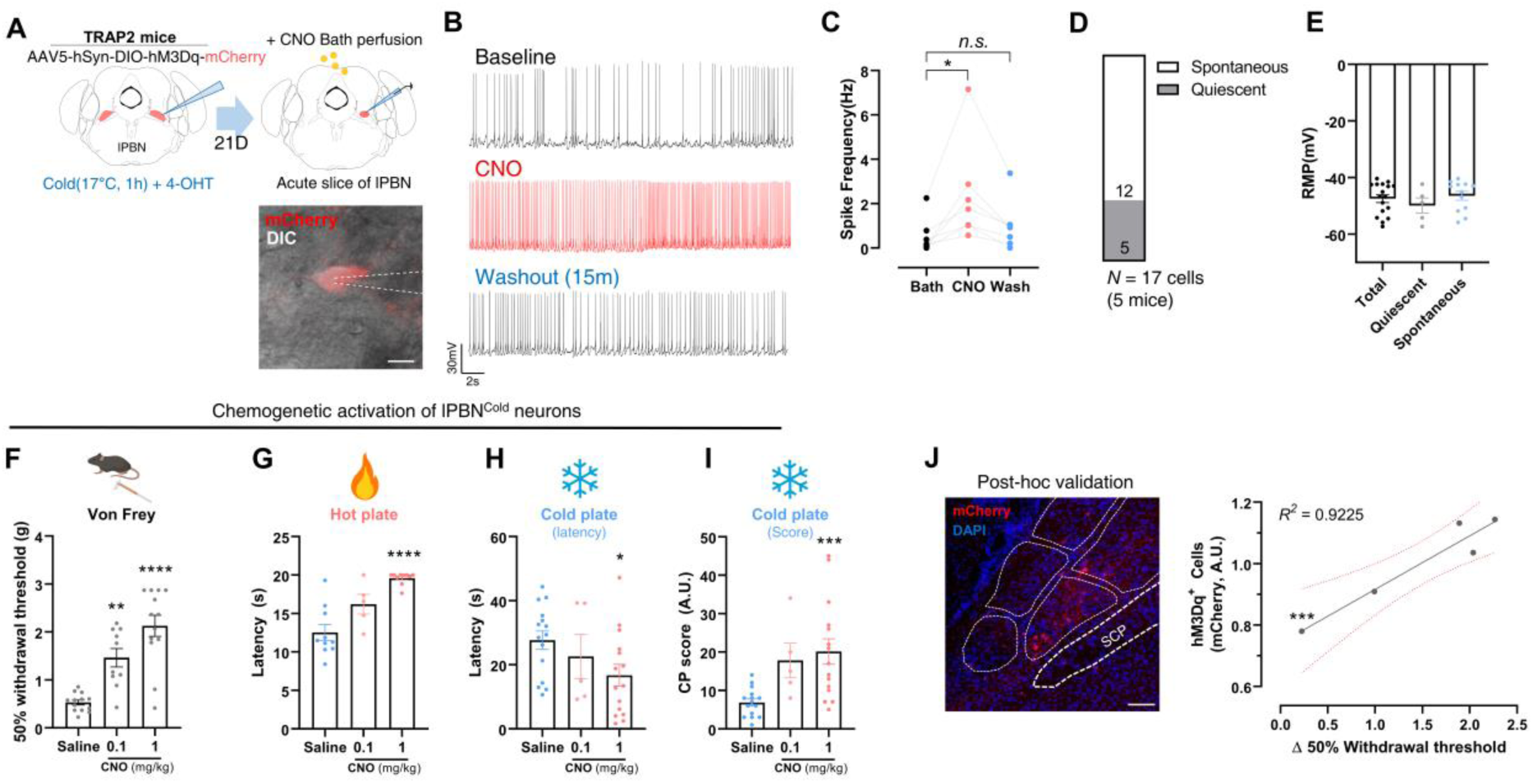
Chemogenetic activation of lPBN^cold^ neuron induces acute antinociception. (A) Experimental scheme for the experiments presented in Figure 3A-E. Red: mCherry. Scale bar: 10μm. (B) Representative membrane potential trace of lPBN^cold^ neurons during CNO perfusion. (C) Quantification of CNO-induced spike frequency in lPBN^cold^ neurons. (D) Proportion of spontaneously firing neurons and quiescent neurons in recorded lPBN^cold^ neurons. (E) Resting membrane potential of recorded lPBN^cold^ neurons. (F) Mechanical sensitivity measured with 50% withdrawal thresholds in lPBN^cold^ neuron activation experiment. (G) Thermal sensitivity to noxious hot measured with response latency to hot plate in lPBN^cold^ neuron activation experiment. (H) Thermal sensitivity to noxious cold measured with response latency to cold plate in lPBN^cold^ neuron activation experiment. (I) Thermal sensitivity to noxious cold measured with behavioral score to cold plate in lPBN^cold^ neuron activation experiment. (J) Representative image of cold-TRAPed lPBN neurons (left) and correlation plot of antinociceptive effect and expression of DREADD in lateral parabrachial nucleus (right). Red: mCherry, Blue: DAPI. Scale bar: 100μm.

### Analgesic role of lPBN^cold^-to-vlPAG neural circuit through opioid receptor signaling in vlPAG

To identify potential downstream targets of lPBN^cold^ neurons that mediates analgesic effects, we performed an anterograde neural tracing on lPBN^cold^ neurons (**Fig. 4A**). Tracing experiments revealed axonal projection of lPBN^cold^ neurons to the ipsilateral PAG, mainly in the ventrolateral subcolumn, along with projections to paraventricular thalamus (PVT) and medial hypothalamus (**Fig. 4B**). The involvement of vlPAG in pain relief is well established, and activation of lPBN to PAG projection induces antinociception according to previous reports (*24*). Moreover, we observed significant activation of vlPAG following cooling stimuli (**Fig. 1E**). Thus, we focused on the connectivity and functional role of lPBN and vlPAG in the following investigation. To functionally validate the role of vlPAG projection of the lPBN^cold^ neurons, we first investigated whether cold stimulation recruits the lPBN-to-vlPAG pathway. Cold stimuli induced Fos expression in 17.8% of vlPAG-projecting lPBN neurons, and 27.8% of cold-positive neurons were projecting to vlPAG (**fig. S7,** See **Table S1**). Given that cold stimuli also significantly increased neuronal activity in the vlPAG (**Fig. 1**) and lPBN-vlPAG neural circuit has been previously implicated in antinociception (*25*), we sought to define the role of lPBN-vlPAG neural circuit in TRPM8-mediated analgesia. We selectively activate vlPAG projection of lPBN^cold^ neurons by CNO microinjection into bilateral vlPAG (**Fig. 4C**). Similar to global lPBN^cold^ neuron activation via intraperitoneal injection of CNO, vlPAG-targeted CNO administration significantly increased mechanical and heat pain threshold, while decreasing response latency to noxious cold, mimicking behavioral phenotype observed following general lPBN^cold^ neuron activation (**Fig. 4D-G**). To rule out potential confounds related to altered motivation or motor function, we conducted a sticky tape removal test applied to paw. Selective activation of the lPBN^cold^-vlPAG projection did not affect the latency to attending for the tape attached at forepaw, nor the latency to remove the tape, indicating that the observed analgesic effects were not attributable to generalized suppression of motor function and/or reduced motivational drive to respond (**Fig. 4H**). Given that the vlPAG is a critical hub for endogenous opioid-mediated analgesia and cold-induced analgesia requires opioid receptor signaling (*40, 41*), we examined whether local opioid receptor signaling in the vlPAG contributes to lPBN^cold^ neuron activation-induced analgesia. To test this hypothesis, we microinjected the opioid receptor antagonist naloxone into the bilateral vlPAG while activating the lPBN^cold^- vlPAG neural circuit. Local naloxone treatment into vlPAG significantly attenuated the mechanical antinociception induced by lPBN^cold^-vlPAG circuit activation, suggesting that opioid receptor signaling in the vlPAG is necessary for lPBN^cold^-vlPAG neural circuit mediated analgesia (**Fig. 4I**). Lastly, chemogenetic activation of lPBN^cold^-vlPAG neural circuit activation was sufficient to reverse CFA-induced mechanical allodynia (**Fig. 4J**). Collectively, our data demonstrated the antinociceptive role of lPBN^cold^- vlPAG neural circuit, which is mediated by local opioid receptor signaling in the vlPAG.

**Figure 4.**
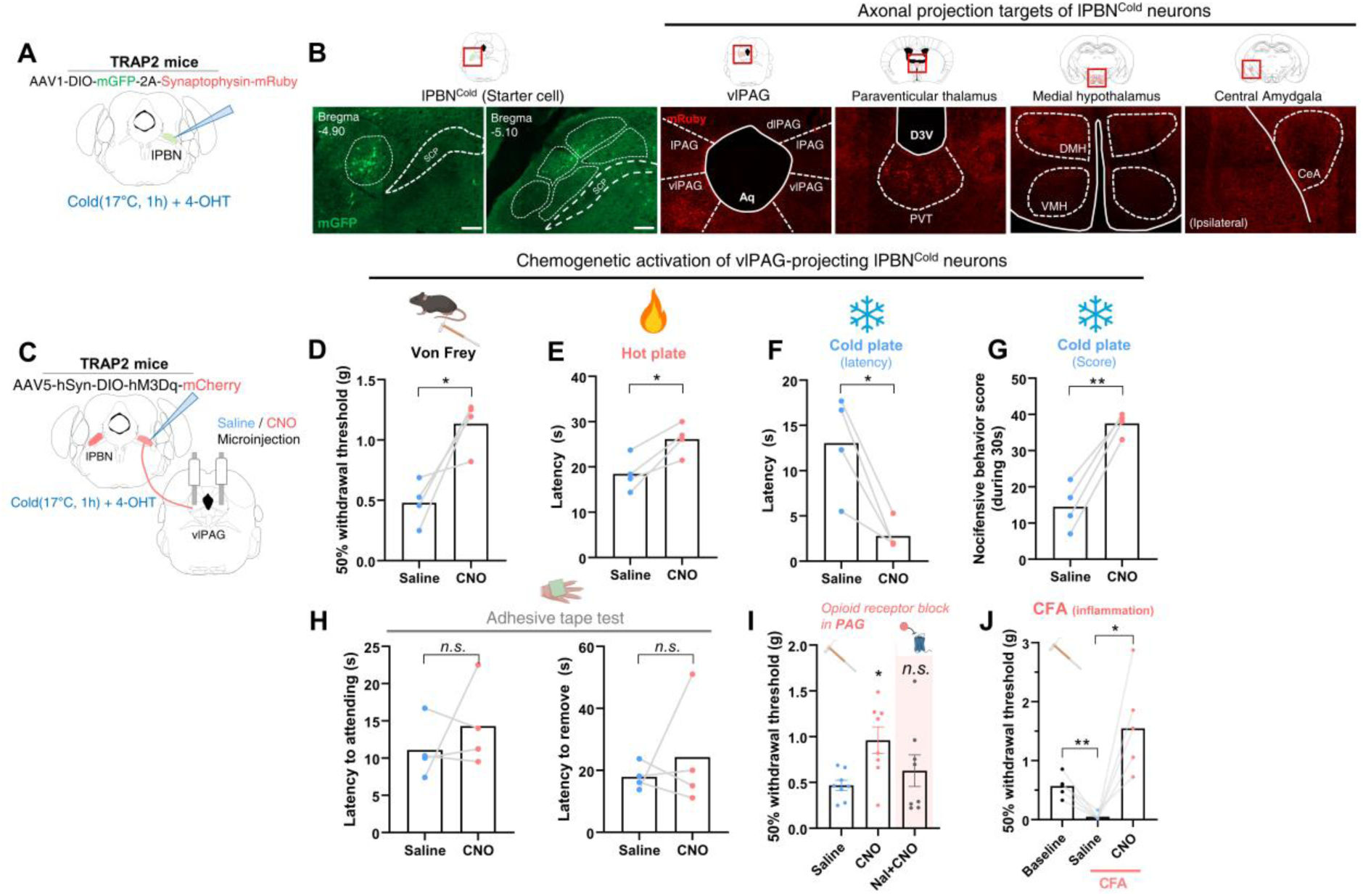
Analgesic role of lPBN^cold^-to-vlPAG neural circuit through opioid receptor signaling in vlPAG. (A) Experimental scheme for the anterograde tracing of lPBN^cold^ neurons. (B) Distribution of GFP-positive (Cre^+^) neurons and neuronal axon projection in a diverse brain region. Green; GFP. Red; Synaptophysin-mRuby. Scale bar: 100μm. (C) Experimental scheme for the selective activation of vlPAG-projecting lPBN^cold^ neurons. (D) Mechanical sensitivity measured with 50% withdrawal thresholds in vlPAG-projecting lPBN^cold^ neuron activation experiment. (E) Thermal sensitivity to noxious hot measured with response latency to hot plate in vlPAG-projecting lPBN^cold^ neuron activation experiment. (F) Thermal sensitivity to noxious cold measured with response latency to cold plate in vlPAG-projecting lPBN^cold^ neuron activation experiment. (G) Thermal sensitivity to noxious cold measured with behavioral score to cold plate in vlPAG-projecting lPBN^cold^ neuron activation experiment. (H) Latency to first attempt to tape removal in vlPAG-projecting lPBN^cold^ neuron activation experiment (left) and latency to tape removal in vlPAG-projecting lPBN^cold^ neuron activation experiment (right). (I) Mechanical sensitivity measured with 50% withdrawal thresholds in vlPAG-projecting lPBN^cold^ neuron activation experiment, followed by local naloxone injection. (J) Mechanical sensitivity measured with 50% withdrawal thresholds in vlPAG-projecting lPBN^cold^ neuron activation experiment, performed in CFA-induced inflammatory pain model mice. Data are presented as mean ± SEMs. ∗*p* < 0.05, ∗∗*p* < 0.01, ∗∗∗*p* < 0.001 and ∗∗∗∗*p* < 0.0001. Number of mice and statistical tests are listed in **Table S1**.

### lPBN^cold^ neurons are necessary for TRPM8-dependent analgesia

Next, we tested whether lPBN^cold^ neurons and their downstream projections are necessary for TRPM8-dependent analgesia. To achieve this, we selectively ablated lPBN^cold^ neurons with Casp3 (**Fig. 5A-C**). Given that previous studies have implicated the lPBN in homeostatic temperature regulation (*18, 42*), we also evaluated the role of lPBN^cold^ neurons in thermoregulation and thermal preference, assessed by preferred temperature in a spontaneous thermal gradient test. Ablation of bilateral lPBN^cold^ neurons did not alter baseline thermal preference and nociceptive sensitivity including mechanical and thermal pain (**Fig. 5D**, **fig. S7A-D, F-H**). Additionally, ablation of lPBN^cold^ neurons did not affect core body temperature in basal state, as measured by rectal temperature probe (**fig. S7E**). To determine the necessity of lPBN^cold^ neurons for TRPM8-mediated analgesia, we employed the same behavioral paradigm described in **Fig 1**. To rule out the potential effect of icilin treatment procedure *per se*, which include light restrain and potential stress, we compared the mechanical withdrawal threshold between baseline and vehicle-treated mice, which revealed no significant differences (**Fig. 5E**). Ablation of the lPBN^cold^ neurons significantly attenuated mechanical antinociception induced by icilin administration to hindpaw in both naïve and CFA-induced inflammatory pain states (**Fig. 5F-H**), demonstrating the essential role of lPBN^cold^ neurons in TRPM8-mediated analgesia. Furthermore, selective chemogenetic inactivation of the lPBN-to-vlPAG neural circuits phenocopied the loss of TRPM8-mediated analgesia, providing additional evidence that the lPBN-to-vlPAG circuit is a critical component of TRPM8-dependent pain modulation (**Fig. 5I, J**).

**Figure 5.**
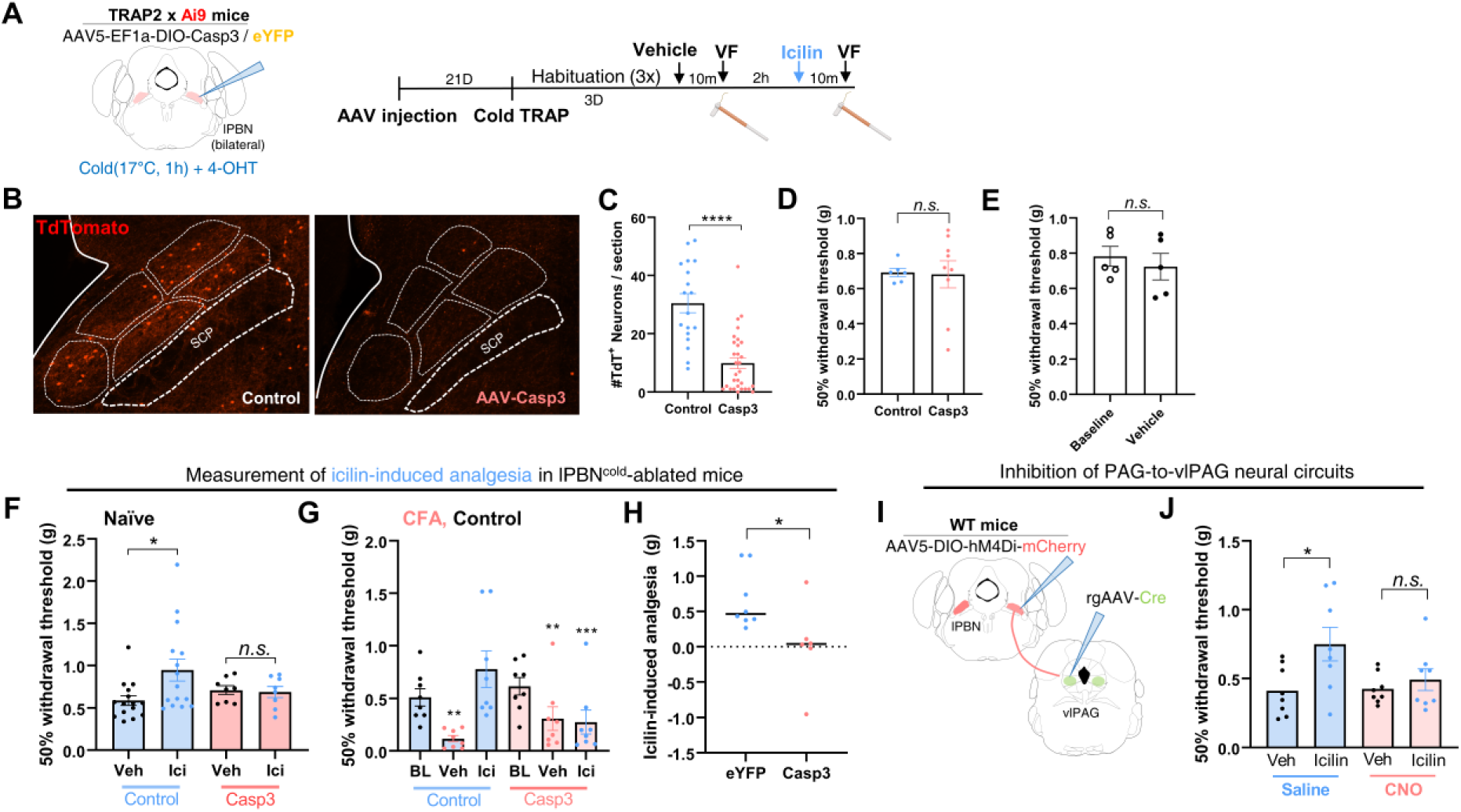
lPBN^cold^ neurons are necessary for TRPM8-dependent analgesia. (A) Experimental scheme and timeline for the icilin-induced pain relief experiment. (B) Representative images of lPBN in control and Casp3-medated lPBN^cold^ neuron ablated mice. Red; TdTomato. (C) Quantification of TdTomato^+^ neurons per lPBN section in control and Casp3-medated lPBN^cold^ neuron ablated mice. (D) Mechanical sensitivity measured with 50% withdrawal thresholds in control and Casp3-medated lPBN^cold^ neuron ablated mice. (E) Mechanical sensitivity measured with 50% withdrawal thresholds in control mice, before and after vehicle treatment. (F) Mechanical sensitivity following icilin treatment, in control mice and casp3-mediated lPBN^cold^ neuron ablated naïve mice. (G) Mechanical sensitivity following icilin treatment, in control mice and casp3-mediated lPBN^cold^ neuron ablated mice after intraplantar CFA injection. (H) Quantification of topical icilin-induced antinociception (measure in difference of 50% withdrawal threshold) in control mice and casp3-mediated lPBN^cold^ neuron ablated mice. (I) Experimental scheme for lPBN-to-vlPAG neuronal circuit inhibition. (J) Mechanical sensitivity following icilin treatment, in saline- and CNO-treated mice for lPBN- to-vlPAG neuron inhibition. Data are presented as mean ± SEMs. ∗*p* < 0.05, ∗∗*p* < 0.01, ∗∗∗*p* < 0.001 and ∗∗∗∗*p* < 0.0001. Number of mice and statistical tests are listed in **Table S1**.

### Identification of enkephalinergic signaling within vlPAG in TRPM8-dependent analgesia via transcriptomic profiling of lPBN^cold^ neurons

The lPBN is a highly heterogeneous structure, comprised of multiple histologically and genetically distinct subnuclei, each with specialized physiological functions (*34*). To genetically characterize lPBN^cold^ neurons, we used RNA sequencing combined with a fluorescence-activated cell sorter (FACS) (See methods; **Fig. 6A**). Prepared single cell suspension was sorted based on green fluorescence (indicative of cold responsiveness) and DAPI (indicates cell integrity and viability) through FACS, yielding a 79.34% of viable cells and 5.88% of Zsg-positive cells among singlet gated counts (**Fig. 6B-D**). To determine differentially expressed genes, we performed RNA sequencing on sorted viable Zsg-positive and Zsg-negative populations. Comparative analysis identified 207 significantly enriched genes and 1036 significantly depleted genes in Zsg-positive cells relative to Zsg-negative cells (**Fig. 6E**). As Zsg-positive TRAP cells were predominantly neurons (*43*), classical markers for microglia (*Cx3cr1, Tmem119*), astrocyte (*Gfap, S100b*) and oligodendrocytes (*Mag, Mbp*) were significantly depleted or at least not enriched in Zsg-positive cells, and general housekeeping genes (*Actb, Gapdh*) remain unchanged between groups. Among the significantly enriched genes, we annotated genes associated with neuropeptide signaling, endogenous opioid pathways and established marker genes for parabrachial nucleus. Notably, *Penk* (proenkephalin), a precursor for enkephalin, was significantly enriched in Zsg-positive cells and RNAscope FISH analysis validated that 46.2% of Cold-TRAP (Zsg-positive) lPBN neurons express RNA transcript for *Penk* (**Fig. 6F**). This suggests a potential molecular basis for cold-induced analgesia via endogenous opioid signaling. Indeed, chemogenetic activation of lPBN^cold^ neurons significantly increased the enkephalin level measured with immunohistochemistry in the vlPAG, suggesting the functional expression and release of enkephalin by lPBN^cold^ neurons (**Fig. 6G-I**). Moreover, exposing hindpaws to cold significantly increased the enkephalin level in vlPAG in naïve, wild-type mice (**Fig. 6J**). Previous report demonstrated that enkephalin signaling within the vlPAG mediates analgesic effect (*44, 45*). Given that local opioid receptor signaling was necessary for lPBN^cold^-vlPAG neural circuit mediated antinociception (**Fig. 4I**), we hypothesized that enkephalin release in the vlPAG represents a potential mechanism underlying TRPM8-mediated analgesia. To directly test this hypothesis, we administered a neutralizing antibody against enkephalin bilaterally into the vlPAG before behavioral testing (**Fig. 6K**). Antibody-mediated enkephalin blockade significantly attenuated the mechanical antinociception induced by icilin administration to hindpaw, compared to saline-injected controls (**Fig. 6L**). These findings demonstrated that enkephalinergic signaling within the vlPAG plays a crucial role in mediating TRPM8-mediated analgesia.

**Figure 6.**
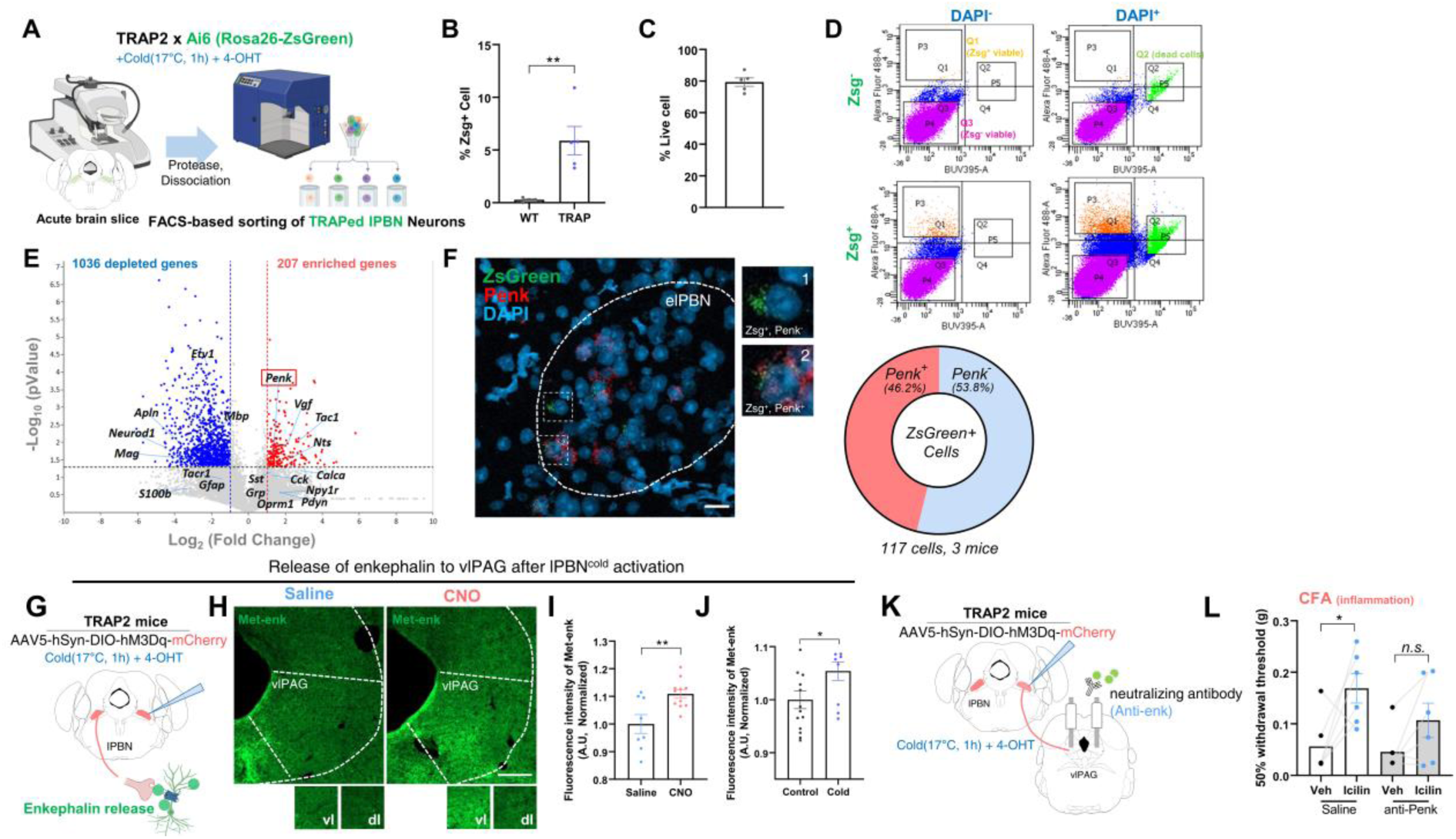
Identification of enkephalinergic signaling within vlPAG in TRPM8-dependent analgesia via transcriptomic profiling of lPBN^cold^ neurons. (A) Experimental scheme for preparation of single-cell suspension and sorting of Zsg-positive neuron of lPBN. (B) Percentage of Zsg-positive cells in total viable cells in Wildtype mice and TRAP2 x Ai6 mice. (C) Percentage of viable cells in total counts. (D) Gating strategy for fluorescence-based sorting of Zsg-positive, viable neuron from brain single cell suspension. (E) Volcano plot of significantly enriched genes (Red dot) and depleted genes (Blue dot) in Zsg-positive cells compared to Zsg-negative cells. (F) Representative figure of RNAscope fluorescence in situ hybridization in lPBN (left) and quantification of marker co-expression in Zsg-expressing lPBN neurons (right). Red; Penk, Green; Zsgreen, Blue; DAPI (left) Scale bar: 20μm. (G) Experimental scheme for the selective activation of lPBN^cold^ neurons for enkephalin release measurement. (H) Representative figure of met-enkephalin immunostaining in vlPAG, with lPBN^cold^ neurons activity manipulation (top) and enlarged images from ventrolateral subcolumn (vl) and dorsolateral subcolumn (dl). Green; Met-enkephalin. Scale bar: 100μm. (I) Quantification of met-enkephalin level in vlPAG after lPBN^cold^ neuron activity manipulation. (J) Quantification of met-enkephalin level in vlPAG, in control mice and cold-exposed mice. (K) Experimental scheme for the local neutralizing antibody treatment in vlPAG before TRPM8-mediated analgesia behavioral tests. (L) Mechanical sensitivity measured with 50% withdrawal thresholds in CFA-induced inflammatory pain model, with neutralizing antibody administration in the vlPAG. Data are presented as mean ± SEMs. ∗*p* < 0.05, ∗∗*p* < 0.01, ∗∗∗*p* < 0.001 and ∗∗∗∗*p* < 0.0001. Number of mice and statistical tests are listed in **Table S1**.

### Activation of lPBN^cold^ neurons decreases Fos expression to noxious stimuli in the spinal cord, with involvement of descending pain modulation circuit

The vlPAG contains opioid receptor-expressing neurons, playing a crucial role in endogenous opioid mediated analgesia and descending pain modulation through their interactions with other brainstem regions such as locus coeruleus (LC) and rostral ventrolateral medular (RVM) (*46, 47*). These structures are known to project to the spinal cord and induce antinociception via noradrenergic or serotonergic signaling, respectively (*48*). Thus, we investigated whether cold stimulation activates key brain regions involved in descending pain modulation, as a downstream pathway from the vlPAG. Indeed, exposure to cold stimuli significantly increased TRAP fluorescence signal in vlPAG and RVM (**Fig. 7A, B**), two most well-defined regions involved in descending pain modulatory circuit (*49*). Notably, within the RVM, nucleus raphe pallidus (RPa), which contains serotonergic neurons (*50*), exhibited robust fluorescence expression. To functionally assess the role of lPBN^cold^ neurons in descending pain modulation, we chemogenetically activated lPBN^cold^ neurons and repeatedly applied noxious von Frey stimulus (1.4g) to bilateral hindpaws. After 90 minutes, noxious mechanical stimulus-induced Fos expression in the lumbar spinal cord (L3-5) was assessed (**Fig. 7C**). As expected, activation of lPBN^cold^ neurons significantly reduced noxious mechanical stimuli-induced Fos in both superficial and deep laminae of the spinal cord (**Fig. 7D-G**). These results indicate that lPBN^cold^ neurons modulate nociception through descending pain modulation, potentially through RVM.

**Figure 7.**
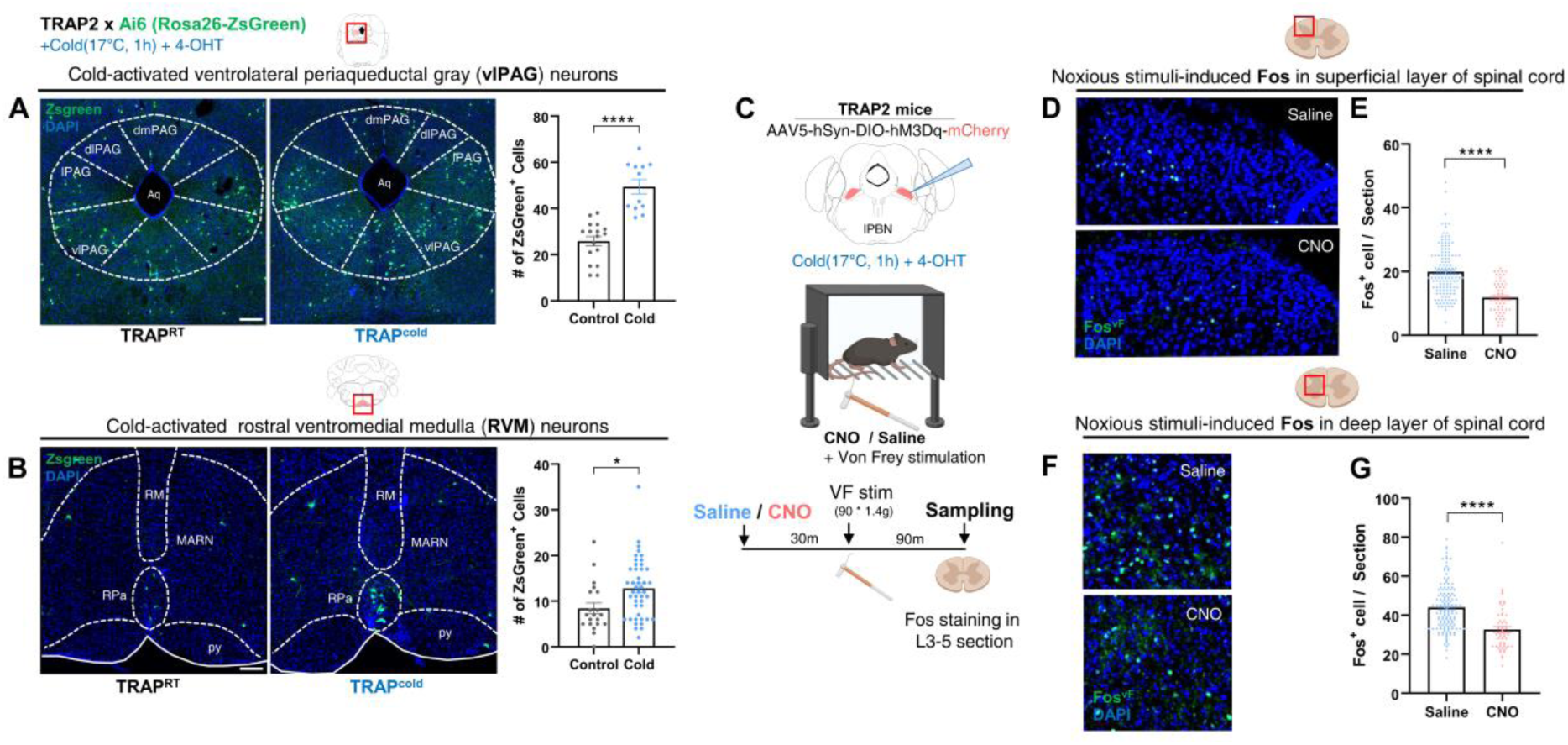
Activation of lPBN^cold^ neurons decreases Fos expression to noxious stimuli in the spinal cord, with involvement of descending pain modulation circuit. (A) Distribution of Zsg-positive neurons in a periaqueductal gray of a control-TRAP mouse and Cold-TRAP mouse (left) and quantification of Zsg-positive neurons in vlPAG (right). Blue; DAPI, Green; Zsgreen. Scale bar: 200μm. (B) Distribution of Zsg-positive neurons in a rostral ventromedial medullar of a control-TRAP mouse and Cold-TRAP mouse (left) and quantification of Zsg-positive neurons in RVM (right). Blue; DAPI, Green; Zsgreen. Scale bar: 200μm. (C) Experimental scheme and timeline for spinal cord Fos immunostaining experiment in lPBN^cold^ neuron activation, presented in Figure 7D-G. (D) Distribution of Fos immunostaining in a superficial layer of lumbar 3-5 spinal cord in a saline- and CNO-treated mice. Blue; DAPI, Green; Fos. (E) Quantification of Fos^+^ Cells in a superficial layer of lumbar 3-5 spinal cord in a saline- and CNO-treated mice. (F) Distribution of Fos immunostaining in a deep layer of lumbar 3-5 spinal cord in a saline- and CNO-treated mice. Blue; DAPI, Green; Fos. (G) Quantification of Fos^+^ Cells in deep layer of lumbar 3-5 spinal cord in a saline- and CNO-treated mice. Data are presented as mean ± SEMs. ∗*p* < 0.05, ∗∗*p* < 0.01, ∗∗∗*p* < 0.001 and ∗∗∗∗*p* < 0.0001. Number of mice and statistical tests are listed in **Table S1**.

### Activation of cold-activated descending pain modulation circuit decrease spinal responsiveness to noxious stimuli

Previous reports have found that neurons reside in RVM contains several types of spinally projecting neurons, categorized based on their responses to noxious stimuli: “on-cells,” “off-cells,” and “neutral cells” (*51*). On-cells are activated by noxious stimulation and are associated with pain facilitation, whereas off-cells are inhibited just prior to nociceptive reflexes and are involved in pain inhibition. Thus, to overcome this functional heterogeneity and directly validate the involvement of descending pain modulation via RVM pain inhibitory neurons, we selectively activated cold-responsive RVM neurons (RVM^cold^ neurons) by expressing excitatory DREADD hM3Dq into RVM^cold^ neurons using TRAP2 transgenic mice (**Fig. 8A-B**). Chemogenetic activation of RVM^cold^ neurons induced significant antinociceptive effect to noxious mechanical and heat stimuli, but not to noxious cold stimuli, demonstrating modality-specific modulation of nociception (**Fig. 8C-E**). Likewise, chemogenetic activation of RVM^cold^ neurons significantly decreased noxious mechanical stimulation-induced Fos in lumbar spinal cord (**Fig. 8F-I**). Collectively, these results suggest the involvement of lPBN^cold^ neurons in the descending pain modulation through antinociceptive RVM neurons, further revealing the neural circuit mechanisms underlying cold-induced analgesia.

**Figure 8.**
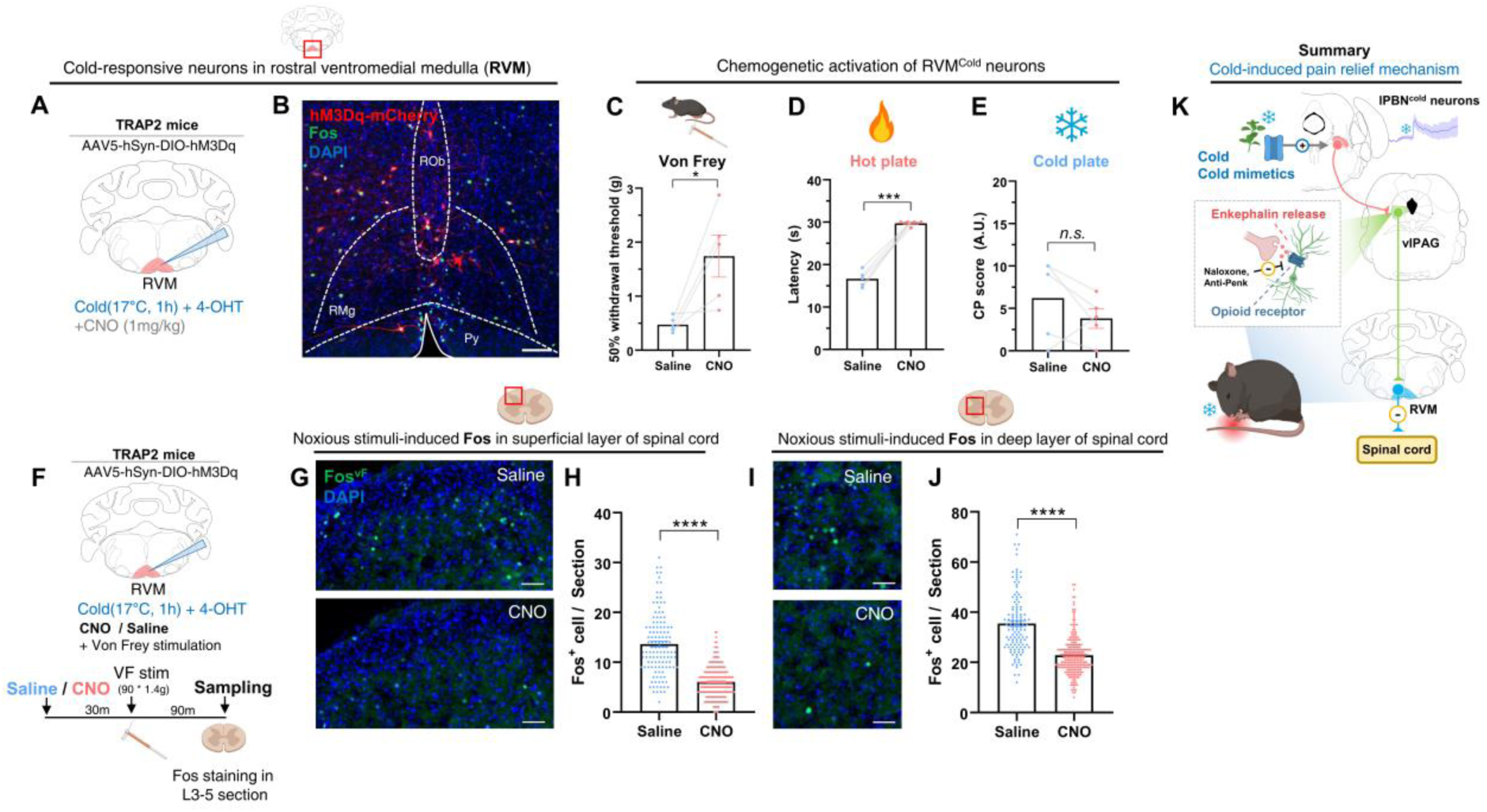
Activation of cold-activated descending pain modulation circuit decrease spinal responsiveness to noxious stimuli. (A) Experimental scheme for the experimental test presented in Figure 8A-E. (B) Representative image of cold-TRAP RVM neurons. Blue; DAPI, Green; Fos, Red; mCherry. Scale bar: 100μm. (C) Mechanical sensitivity measured with 50% withdrawal thresholds in RVM^cold^ neuron activation experiment. (D) Thermal sensitivity to noxious hot measured with response latency to hot plate in RVM^cold^ neuron activation experiment. (E) Thermal sensitivity to noxious cold measured with nocifensive behavioral score to cold plate in RVM^cold^ neuron activation experiment. (F) Experimental scheme for the experimental test presented in Figure 8G-J. (G) Distribution of Fos immunostaining in a superficial layer of lumbar 3-5 spinal cord in a saline- and CNO-treated mice. Blue; DAPI, Green; Fos. Scale bar: 50μm. (H) Quantification of Fos^+^ Cells in a superficial layer of lumbar 3-5 spinal cord in a saline- and CNO-treated mice. (I) Distribution of Fos immunostaining in a deep layer of lumbar 3-5 spinal cord in a saline-and CNO-treated mice. Blue; DAPI, Green; Fos. Scale bar: 50μm. (J) Quantification of Fos^+^ Cells in deep layer of lumbar 3-5 spinal cord in a saline- and CNO-treated mice. (K) Schematic diagram for the mechanism of cold-induced pain relief. Data are presented as mean ± SEMs. ∗*p* < 0.05, ∗∗*p* < 0.01, ∗∗∗*p* < 0.001 and ∗∗∗∗*p* < 0.0001. Number of mice and statistical tests are listed in **Table S1**.

## Discussion

In this study, we first demonstrated that cold-responsive neurons within the lateral parabrachial nucleus governs TRPM8-dependent analgesia through projections to vlPAG by using viral tracing and chemogenetic activity modulation. In contrast, lPBN^cold^ neurons were indispensable for thermal preference and body temperature regulation. Moreover, through transcriptomic analysis combined with TRAP method, we discovered that cold-responsive lPBN neurons were enriched with *Penk*, and TRPM8-dependent pain relief is mediated by local enkephalinergic signaling in the vlPAG. Lastly, activation of cold-responsive lPBN neurons diminished spinal responsiveness to noxious stimuli potentially through cold-responsive RVM neurons, revealing the cellular and molecular level mechanism underlying the cold-induced pain relief (**Fig. 8K**).

The lPBN comprises functionally divergent neuronal populations that process a wide range of sensory stimuli. While previous literatures reported that distinct subpopulation of lPBN neurons are responsive to diverse sensory stimuli such as pruritic (itch-inducing) stimuli, painful pinch and noxious heat, temperature change, gustatory stimuli and mechanical distension from stomach (*17, 52–56*), direct single-cell resolution observation of lPBN neuronal response to peripheral sensory modality has been limited. In this study, using *in vivo* calcium imaging, we identified distinct neuronal populations within the lPBN such that neurons responding to cold are separated from neurons responsive to noxious heat and pinch (**Fig. 2**). This observation aligns with prior extracellular unit recordings in lPBN, which identified cold-sensitive, but not responsive to noxious heat and mechanical pinch stimuli neurons without somatotopic arrangement (*36*). Additionally, *in vivo* calcium imaging of spinoparabrachial (SPB) neurons in the spinal dorsal horn and whole-cell patch clamp recording of SPB neurons with skin-nerve preparation showed the similar responsive pattern, which a subset of SPB neurons are selectively activated by cold, but not to mechanical indentation and noxious heat (*57, 58*). Given that SPB neurons relays primary sensory input from periphery to lPBN, it is likely that the sensory responsiveness pattern of lPBN neurons is largely driven by SPB neurons.

Previous studies have identified cold-responsive neurons within the lPBN primarily in the context of a homeostatic thermoregulation and thermal preference, as activation of DMH-projecting lPBN neurons induced thermogenesis, and ablation of lPBN impaired normal thermal preference (*17, 18, 42*). In contrast, in the present study, neither activation nor inactivation of lPBN^cold^ neurons altered core body temperature (**fig S6F, 8E**) and ablation of lPBN^cold^ neurons did not change thermal preference (**fig. S6**). Instead, activation of lPBN^cold^ neurons modulate acute nociceptive responsiveness (**Fig. 3F-I**), and anterograde tracing of lPBN^cold^ neurons showed relatively sparse DMH projection (**Fig. 4B**). Moreover, RNA sequencing analysis showed lack of somatostatin (SST) enrichment in lPBN^cold^ neurons (**Fig. 5E**). These physiological, anatomical, transcriptomic and behavioral evidences suggest that the lPBN^cold^ neuronal population identified in the current study is distinct from DMH-projecting, SST-positive lPBN neurons which are implicated in thermoregulation in a previous report (*18*). This discrepancy may stem from differences in the stimuli used to label cold-responsive neurons. In our study, cold-responsive neurons were identified and manipulated following relatively short exposure to innocuous cold stimulation with normal ambient temperature, primarily targeting the paw and tail regions via placement on the metal plate (**Fig. 1, fig. S1; see Methods**), while aforementioned studies exposed the mouse to the cold chamber for longer time period, which would challenge and compromise core body temperature (*17, 18*). Accordingly, it is plausible that neurons responsive to core body temperature change that governs long-term, homeostatic thermoregulation are functionally distinguished from those involved in short-term, acute behavioral response regulation to sensory stimuli. Nonetheless, the precise mechanisms by which these neural populations encode different sensory stimuli and regulate distinct physiological functions require further investigation.

In the present study, we identified a subset of lPBN neurons projecting to the ventrolateral periaqueductal gray (vlPAG) that exert antinociceptive effects (**Fig. 4D,E**). Notably, the axonal projection of lPBN^cold^ neurons was predominantly localized in the vlPAG, whereas prior tracing studies have shown that noxious stimulus-responsive TacR1^+^ lPBN neurons primarily target the dorsolateral division of PAG (dlPAG), a region implicated in active flight and nocifensive response (*34, 53*). Furthermore, we found that majority of these cold-responsive lPBN neurons are *Penk*^+^ neurons, suggesting the they constitute a previously uncharacterized subpopulation of lPBN neurons that mediate analgesic action, potentially through enkephalin release within the vlPAG (**Fig. 6G-J**). The vlPAG is one of the most well-characterized brain regions involved in opioid-mediated analgesia, including enkephalin-mediated analgesia (*45, 59–62*). For instance, previous studies have demonstrated that microinjection of enkephalin or enkephalinase inhibitor into vlPAG is antinociceptive (*63, 64*). Moreover, administration of anti-enkephalin neutralizing antibody into vlPAG diminishes analgesic effect of electroacupuncture, highlighting the functional role of enkephalin signaling in vlPAG in pain suppression (*44*). To note, conventional tracing and anatomical studies have confirmed the presence of enkephalinergic neuronal terminals in vlPAG, yet the precise source of enkephalinergic neuronal terminals has remained elusive (*65*). Recent evidence has shown that the lPBN contains enkephalinergic neurons that project to the vlPAG (*45*), which aligns with current experimental results. Moreover, our study revealed that activation of lPBN^cold^ neurons and cold exposure significantly increased local enkephalin levels in the vlPAG (**Fig. 6I,J**), further supporting the functional role of lPBN as a source of enkephalin in the vlPAG. Especially, majority of lPBN^cold^ neurons displayed spontaneous activity (**Fig. 3B,C, fig S5**), which could maintain the tonic release of enkephalin in PAG (*66*). Previous studies have consistently demonstrated that cold-induced analgesia relies on opioid receptor signaling, but the direct action of this signaling has remained elusive (*1, 4, 13*). Bridging a knowledge gap, we first demonstrated that enkephalinergic signaling in the vlPAG contributes to TRPM8-induced analgesia (**Fig. 4I**, **Fig. 6L**), via RVM-mediated descending pain modulation pathway (**Fig. 7**, **Fig. 8**). Supporting our experimental data, neuroimaging studies conducted in human subjects have demonstrated that analgesic cold stimuli significantly activate the PAG (*48, 67, 68*), and pain relief induced by skin cooling and menthol is significantly attenuated by the opioid receptor antagonist naloxone (*1, 4, 13*). This suggested that cold-induced analgesia could also be mediated in human through central pathway involving the PAG, enkephalinergic opioid receptor signaling and the descending pain inhibitory circuit.

The aforementioned experimental results suggest that cold-induced analgesia is primarily mediated by the central nervous system, particularly supraspinal structures but not peripheral level (**fig S2**). However, others demonstrated that formalin-induced spontaneous calcium response in dorsal root ganglion can be inhibited by topical menthol application and physical cooling inhibits spontaneous calcium responses in injured trigeminal ganglia neurons (*69, 70*). This might contribute to decreased activity of lPBN^HP^ neurons upon cold exposure (**Fig. 2G**). Moreover, recent studies have shown that activation of cold-responsive Kcnip2^+^ neurons in the spinal cold could induced analgesia (*71*), suggesting a possible intraspinal local inhibitory processing in TRPM8-dependent analgesia. Indeed, previous research has reported that intrathecal injection of a Group II/III mGluR inhibitor, but not naloxone, abolished TRPM8-dependent analgesia (*28*). These body of evidence suggests that TRPM8-dependent analgesia may be mediated at multiple levels, from peripheral nervous system, spinal cord and brainstem.

In conclusion, our study elucidates the role of lPBN-to-vlPAG neural circuit and local enkephalin signaling in TRPM8-dependent analgesia, highlighting a potential central target for novel pain therapeutics.

## Resource availability

Further information and requests for resources and reagents should be directed to the lead contact, **Seog Bae Oh**, DDS, PhD.

## Materials availability

This study did not generate new, unique reagents.

## Data and code availability

All data reported in this paper will be shared by the lead contact upon request; This paper does not report any original code; Any additional information required to reanalyze the data reported in this paper is available from the lead contact upon request.

## Acknowledgements

This research was supported by the National Research Foundation of Korea (NRF) grant funded by the Ministry of Science and ICT, Republic of Korea (grant number: RS-2021-NR059709, RS-2023-00264409, RS-2024-00441103).

## Author contributions

**Hayun Kim**: Conceptualization, Methodology, Software, Formal analysis, Investigation, Writing-Original Draft, Visualization; **Yoonkyung Lee**: Methodology, Validation, Formal analysis, Investigation, Writing-Original Draft, Visualization; **Seog Bae oh**: Conceptualization, Resources, Data Curation, Writing-Original Draft, Supervision, Project administration, Funding acquisition.

## Methods

### Animals

All behavioral studies and primary culture except the TRAP studies were performed with 5- to 7-week-old male C57BL/6J wild-type (WT) mice (Doo Yeol Biotech, South Korea). For TRAP experiments, 6- to 8-week-old male TRAP2 mice (Fos^2A-iCreER^, JAX strain #030323) were used, crossed with reporter mice (Ai6, JAX strain #007906 or Ai9, JAX strain #007909). They were backcrossed with C57BL/6J mice for more than 10 generations and were thought of as congenic to C57BL/6J. All animals were maintained in constant temperature (22 ± 1°C), humidity (55%), and 12-12h light/dark cycle environment with standard lab chow and water available *ad libitum*. Handling and care of the animals were in accordance with the Guideline for Animal Experiments, 2000, edited by the Korean Academy of Medical Sciences, which is consistent with the National Institutes of Health (NIH) Guidelines for the Care and Use of Laboratory Animals, revised 1996. All experimental procedures were approved by the Institutional Animal Care and Use Committee (IACUC) at Seoul National University (protocol code: SNU-241105-5).

### Primary culture or dorsal root ganglion (DRG) neurons

Primary culture of DRG neuron was performed as previously reported (*72*). Briefly, after the mouse was euthanatized with isoflurane, DRG were acutely dissected on ice-cold HBSS with 20 mM HEPES (Gibco #14025092). Then, ganglia were digested for 1 hour in collagenase A (1 mg/ml, Roche) and dispase II (2.4 U/ml, Roche) mixture in HBSS-HEPES-buffered solution at a 37°C incubator with 5% CO_2_. Then, 0.25% trypsin (Gibco #15090046) in HBSS was applied for 7 minutes. DRG were mechanically dissociated with trituration by fire-polished Pasteur pipette in Dulbecco’s modified Eagle’s medium (DMEM) containing DNase I. The cell suspension was then carefully layered over on 15% bovine serum albumin (BSA) in Ham’s F-12 (Welgene #LM010-52) to isolate neurons. Cells were resuspended with neurobasal medium (Gibco #21103049) containing B-27 supplement (Gibco #17504044), L-glutamine (Sigma-Aldrich #G7513), and penicillin-streptomycin (Gibco #10348016). 3000 DRG neurons were plated on a 10 mm-diameter coverslip coated with poly-D-lysine (100 μg/ml) for calcium imaging experiments.

### Ratiometric calcium imaging in primary sensory neurons

Cultured DRG neurons or transfected HEK-293 cells were used for calcium imaging after incubation overnight. Cells were loaded with 2 μM Fura-2 AM in DMEM for 40 minutes at a 37°C incubator with 5% CO_2_. Images were acquired through a CCD camera (Andor Technology, DL-604M-VP) coupled to an inverted microscope (Nikon, Eclipse TE-2000) while an ultraviolet light source (Sutter Instruments, Lambda DG-4 plus) was used to excite Fura-2 by alternating 340 nm and 380 nm wavelength lights. Mean fluorescence intensity ratios (F340/F380) were measured with MetaFluor software. A bath solution (2 mM Ca^2+^-HBSS containing (in mM) 140 NaCl, 5 KCl, 1 MgCl_2_, 2 CaCl_2_, 10 HEPES, and 10 glucose) was adjusted to pH 7.4 with 5N NaOH and to 300 ± 5 mOsm.

### Behavioral test

#### von Frey test

Before the behavior test, mice were habituated for at least 3 consecutive days to the test chamber and randomized for treatment. All sensory testing was performed between the hours of 09:00 and 18:00 in an isolated room maintained at 22 ± 2°C and 50 ± 10% humidity. For mechanical threshold (von Frey filament) testing, mice were brought from the home cage and placed in transparent plastic boxes on a metal mesh floor. The mice were then habituated for at least 30 min prior to testing. To assess mechanical sensitivity, the withdrawal threshold of the affected hind paw was measured using a series of von Frey filaments (0.20, 0.40, 0.70, 1.6, 3.9, 5.9, 9.8 and 13.7 mN, Stoelting, Wood Dale, IL, USA; equivalent in grams to 0.02, 0.04, 0.07, 0.16, 0.40, 0.60, 1.0 and 1.4). The 50% withdrawal threshold was determined using the ‘up-down’ method. A brisk hind paw lift, licking or flinch in response to von Frey filament stimulation was regarded as a withdrawal response. The 0.4 g filament was the first stimulus to be used, and, when a withdrawal response was obtained, the filament with a next lower number was used. This process was repeated until no response was obtained, at which time the filament with a next higher was administered. All behavioral testing was performed by a blinded investigator who was blind to the treatment of the mice. For complete Freund’s adjuvant (CFA)-induced inflammation model, 20 µl CFA (Sigma, #F5881) was injected in the ipsilateral hind paw 1 days prior to the behavioral test.

#### Drug treatment

CNO was stored as stock solution (10mg/mL in saline, Tocris, #4936) and stored at - 20°C. Before CNO-induced chemogenetic neuronal activation and/or inactivation experiments, stock solution was thawed and diluted to desired concentration with sterile saline. For intraperitoneal injection, chemogenetic activation dose was set as 1mg/kg unless otherwise noted and CNO solution was given with a volume of 10μL/g body weight. Chemogenetic inactivation dose was set as 5mg/kg unless otherwise noted and CNO solution was given with a volume of 10μL/g body weight. For cannula-based local injection of CNO, 1mM CNO dissolved in saline was used. Under light isoflurane anesthesia (1.5%, within 5 minutes), 500nL of CNO solution was filled into microsyringe (Trajan Scientific, #002050) and injected through guide cannula with syringe pump (KdScientific, #KDS100) with a rate of 500nL/minute. Syringe needle tip was remained at place for at least 1 minutes before removal to prevent leakage. CNO injection was performed at least 15 minutes before behavioral test, considering pharmacokinetics of CNO in central nervous system. CNO treatment was randomly counterbalanced with saline to prevent any potential bias based on treatment order. For neutralizing antibody injection into vlPAG, 500nL of 1:1000 anti-met-enkephalin (Immunostar, #20065) diluted in sterile saline solution was administered with a rate of 500nL/min through guide cannula, under light isoflurane anesthesia (1.5%, within 5 minutes). Syringe needle tip was remained at place for at least 1 minutes before removal to prevent leakage. Neutralizing antibody injection was performed at least 15 minutes before behavioral test. For naloxone-mediated local block experiments, 5μg of naloxone (Tocris, #0599) was co-injected with CNO in a volume of 500nL.

#### Topical icilin treatment and cooling-induced analgesia

Icilin (Tocris, #1531) was stored as stock solution of 10mM in DMSO and stored at - 20°C. On the day of treatment, icilin was diluted to vehicle (2% Tween-80 in sterile saline) at a concentration of 200μM. For topical icilin treatment, mice were lightly restrained and icilin solution (or vehicle for control) was applied to bilateral hindpaw with cotton tips for twice, each in 1 minutes. For icilin treatment in tail, tail of mice was emerged into icilin solution twice, each in 1 minutes. For cooling-induced analgesia, mice were place on wet metal plate set for 17°C (for cold-treatment group) or 25°C (for RT-treatment group) for 5 minutes before behavioral test. After cold exposure, moist from hindpaw were gently removed and behavior test was performed within 10 minutes after cold exposure.

#### Hot and cold plate assay

To measure noxious thermal pain threshold, mice were placed onto metallic hot plate or custom-made cold plate within an acrylic container. The temperature of hot plate was set at 55°C, while cold plate was set at -5°C. For hot plate, latency to nocifensive behavior (hindpaw licking, flinching and jumping) was measured three times and averaged. Cut-off time of 30 second was used to prevent permanent tissue damage. For cold plate, latency to nocifensive behavior (hindpaw licking, flinching and jumping) was measured two times and averaged. Cut-off time of 60 second was used to prevent permanent tissue damage. For nocifensive score measurement for cold plate, mice were placed onto cold plate and nocifensive behaviors were counted for 30 second. Every bout of hindpaw licking, flinching and jumping behavior was scored 2 points, and every bout of rearing, hindpaw lifting and trial for chamber avoidance (digging-like behavior) was scored 1 point.

#### Thermal gradient test

The cooling and heating devices were used to maintain stable temperature so that the surface temperature range of the apparatus was from 5°C to 55 °C, in a linear manner. Floor surface temperature was monitored using an attached thermometer. Mice were acclimated for 30 min in the thermal gradient apparatus with its floor at room temperature before the day of the thermal gradient test. A camera is located upon the apparatus to record position of mice in real-time (60 frames per second). Mice were placed individually in the device in an innocuous middle zone and the spontaneous movement of mice was videotaped for 30 minutes. Recorded video was analyzed with ezTrack to measure travel distance or spent time in each region (or Temperature Zone) (*73*). Noxious cold zone was defined as zone with a temperature of 17°C or lower; Cool zone was defined as zone with a temperature between 17°C and 25°C; noxious hot zone was defined as zone with a temperature of 42°C or higher. “preference temperature” was calculated with the average value using the zone temperature and “spent time”.

#### Adhesive tape removal test

Adhesive tape removal test was performed as previously described (*74*). Briefly, after habituation of the animal to the test chamber, patch of adhesive tape (3mm x 4mm) was attached at a central hairless part of the left forepaw. After tape placement, animal was placed in the test chamber and latency to tape contact and tape removal was measured with a timer. The contact time is defined as the point that the mouse reacts to the presence of the adhesive tape strips, with its paw or mouth. Experiments were repeated two times with at least 10 minutes of interval, then the latency obtained from two tests was averaged. If the animal urinates and walks on its micturition before attempting to remove the adhesive tape, session was immediately aborted, and subsequent trial was performed after 10 minutes.

#### Core body temperature measurement

Under light isoflurane anesthesia (1.5%, within 5 minutes), rectal probe covered with vaseline was slowly and carefully inserted to the rectum of animal to prevent damage. After 30 seconds of thermal balancing, core body temperature was measured for 1 minutes and averaged. To measure core body temperature of animals with CNO-induced activation of lPBN^Cold^ neurons, CNO was treated 15 minutes before measurement. Control animals were given same dose of saline. CNO and saline treatment was counterbalanced.

#### 4-OHT treatment

Homozygote TRAP2 mice (Fos2A-iCreER, Jackson #030323) or TRAP2 mice crossed with Ai6 mice (RCL-ZsGreen, Jackson #007906) were used for experiments. 3-4 weeks after stereotaxic AAV injection (in case of AAV injection) or at age of 6-8 weeks (in case of TRAP2; Ai6 mice), mice were handled and treated with saline for acclimation for 5 consecutive days. On the day of treatment, mice were placed on 17°C metal plate powered with peltier devices for 1 hour for Cold TRAP, or left at Room temperature for control TRAP. Immediately after cold exposure, 4-OHT (20mg/kg, dissolved in 2% Tween-80 and 5% DMSO in saline, Sigma, #H6278) was intraperitoneally injected.

#### Stereotaxic surgery and cannula implant

During isoflurane anesthesia (1.5-2%), disinfected scale of mice was cut opened after shaving. Drop of hydrogen peroxide was applied to remove periosteum, then 0.5mm burr hole was made with hand drill. Glass micropipette pulled from glass capillary with Digital puller was filled with mineral oil and AAV solution (>10^13^ vg/mL, Addgene, list attached below) at the end. After slow insertion of micropipette, AAV solution was injected at lPBN (AP -5.00, ML ±1.27, DV -3.27; mm from Bregma), vlPAG (AP -4.60, ML ±0.50, DV -2.60; mm from Bregma) or RVM (AP -5.80, ML 0.00, DV -6.00; mm from Bregma) at a rate of 50nL/min with WPI microinjection (WPI, Nanoliter-2020). Micropipette was withdrawn after 5-10 minutes. Scalp was sutured, then mice are allowed to recover and gene expression for >3 weeks. For GRIN lens insertion surgery for *in vivo* calcium imaging, 400 μm-diameter optic cannula (RWD, #R-FOC-L400C-50NA) was slowly inserted to 250μm above the lPBN. 30 minutes after insertion, cannula was removed and GRIN lens was inserted slowly at a speed of 200 μm/min, using custom holder connected to vacuum was attached to stereotaxic arm to hold GRIN lens (inscopix, #1050-004597) in place. After complete insertion, GRIN lens was attached to skull surface with dental cement and bone wax. To improve adhesion of dental cement to skull, skull surface was scored with scalpel and chemically etched with phosphoric acid gel for 30 seconds and completely dried before cement application. Guide cannula (26G, bilateral 6mm cannula, Protech, #C235-1.0) to vlPAG (AP -4.60, ML ±0.50, DV -2.60; mm from Bregma) was implanted with the same protocol.

|  |  |  |
| --- | --- | --- |
| AAV5-hSyn-DIO-hM3D(Gq)-mCherry | Addgene | Addgene #44361 |
| AAV1-hSyn-DIO-hM4D(Gi)-mCherry | Addgene | Addgene #44362 |
| AAV1-hSyn1-GCaMP6f-P2A-nls-dTomato | Addgene | Addgene #51085 |
| AAV1-hSyn-FLEX-mGFP-2A-Synaptophysin-mRuby | Addgene | Addgene #71760 |
| AAVrg-hSyn-Cre-WPRE-hGH | Addgene | Addgene #105553 |
| AAV5-hSyn-DIO-mCherry | Addgene | Addgene #50459 |
| AAV5-flex-taCasp3-TEVp | Addgene | Addgene #45580 |

#### *In vivo* calcium imaging

After GRIN lens implantation, custom holder was used to mount UCLA miniscope (V4.4) to stereotaxic holder. Miniscope was placed with the right imaging window and baseplate was attached with the dental cement and cured for 30 minutes. After the complete curing, miniscope was removed and protective cap was inserted. Mice were injected with 1mg/kg Dexamethasone to prevent inflammation and swelling for 5 consecutive days after lens implantation and baseplating surgery. After 5 days of Dexamethasone treatment, GRIN lens surface was cleaned with lens paper socked with acetone and ethanol, then imaging window was checked again. If a clear image of cells could not be observed caused by sustained inflammation in imaging windows or distortion of dental cement during curing was observed, mice were excluded from the further functional imaging. For calcium imaging, mice were rightly anesthetized with isoflurane (1.5%) and miniscope was mounted and focused. After full recovery from anesthesia, mice were lightly restrained with head bar and tail of mice were exposed to hot (55°C water), mechanical pinch and cold (4°C water). Cellular response was recorded at a frame rate of 20 frames/second, and raw video was analyzed with Minian pipeline (*75*). Vignetting corrected, denoised, background removed, and motion corrected video was undergone Constrained Nonnegative Matrix Factorization (CNMF) algorithm to estimate temporal and spatial matrix. Finally, cells were manually assessed to exclude any artifacts. Single cells were considered as responsive to stimulus if normalized calcium Z-score was elevated during given peripheral stimulus compared to baseline, and response of neurons to given stimulus in peri-event window was measured with time window of baseline (5 seconds before stimulus) and stimulus (10 seconds after stimulus).

#### Immunohistochemistry (IHC)

For tissue preparation, mice were undergone trans-cardiac perfusion with ice-cold Phosphate-buffered saline (PBS) and 4% paraformaldehyde in PBS. For cold-induced met-enkephalin level measure in vlPAG, mice were perfused and fixed with 4% paraformaldehyde in PBS within 5 minutes after cold stimulation. The harvested brain tissues were fixed overnight in 4% paraformaldehyde in PBS and cryopreserved with 30% sucrose in PBS for at least 48h. Brain and spinal cords were embedded in OCT compound, frozen and cryosectioned by 30 µm per section with cryotome. Brainstem and spinal cord sections were blocked and permeabilized with 4% Normal horse serum with PBS containing 0.3% Triton-X. Sections were washed with PBS, 3 times with 5 minutes each, then stained with 1^st^ antibodies overnight at RT, washed as mentioned above and stained with 2^nd^ antibodies for 2h at RT. All sections were examined and counterstained with DAPI (1μg/mL in PBS) for 15 minutes, washed, then mounted on slide with mounting medium, covered with coverslip and sealed with nail polish. Slides were imaged with Axioscan Slide scanner and analyzed with Zen blue edition and ImageJ. Antibodies used in immunostaining is listed below.

|  |  |  |
| --- | --- | --- |
| Rabbit anti c-Fos | AbCam | #ab190289 |
| Rabbit anti met-enkephalin | ImmunoStar | #20065 |
| Goat anti CGRP | AbCam | #ab36001 |
| Donkey anti rabbit-488 | Jackson ImmunoResearch | #711-545-152 |
| Donkey anti Goat-488 | Jackson ImmunoResearch | #705-545-003 |

#### Preparation of acute brain slices for single cell suspension

Live, single-cell suspensions of brain slices were prepared as described, with a minor modification (*34, 76*). 4-5 male TRAP2; Ai6 mice undergone cold-TRAP procedures and 1 control wild-type mice were used for each experimental batch. All mice were anesthetized with 50mg/kg sodium pentobarbital in saline and perfused with an ice-cold artificial cerebrospinal fluid (aCSF; contraining (in mM) 92 NMDG, 2.5 KCl, 1.25 NaH_2_PO_4_, 30 NaHCO_3_, 20 HEPES, 25 glucose, 2 thiourea, 5 Na-ascorbate, 3 Na-pyruvate, 0.5 CaCl_2_·2H_2_O, and 10 MgSO_4_·7H_2_O. pH was titrated to 7.3–7.4 with hydrochloric acid and solution was continuously bubbled with 95% O_2_ with 95% CO_2_ for at least 20 minutes before use). Brains were rapidly dissected, rested in ice-cold aCSF for 2 minutes, cut, mounted to vibratome with superbond and coronal slices with thickness of 350 µm were prepared using a vibratome from bregma -4.9mm to -5.4mm. Slices were recovered in aCSF containing 500-nM TTX, 10-µM AP-V, and 10-µM DNQX for 30 minutes and lPBN was removed with tissue punches and digested with 1mg/mL Pronase for 50 minutes at 30°C with routine inverting and mechanically dissociated with a pasture pipette (inner diameter 250μm) in a aCSF containing 0.05% BSA. Cell suspensions were filtered through cell strainer tube with a pore size of 35μm twice in 16°C centrifuge (660g, 3 minutes). Cell pellets were suspended with aCSF containing 0.05% BSA and 1μg/mL for 15 minutes for DAPI staining and washed with aCSF containing 0.05% BSA. Remaining cell pellets were resuspended in aCSF containing 0.05% BSA for FACS sorting.

#### Preparation of acute brain slices and electrophysiology

For acute brain slice preparation for ex vivo patch clamp recording, mice were briefly anesthetized with isoflurane and decapitated. Brains were rapidly dissected, rested in ice-cold aCSF for 2 minutes, cut, mounted to vibratome with superbond and coronal slices with thickness of 250 µm were prepared using a vibratome, in ice-cold sucrose-based dissection buffer (containing (in mM) 212.7 Sucrose, 26 NaHCO_3_, 10 glucose, 5 KCl, 10 MgCl_2_, 0.5 CaCl_2_, 1.23 NaH_2_PO_4_; pH was titrated to 7.25 with continuously bubbling with 95% O_2_ with 95% CO_2_ for at least 20 minutes and osmolality was maintained at 330mOsm). Slices were recovered in 28°C artificial cerebrospinal fluid (aCSF; containing (in mM) 124 NaCl, 26 NaHCO_3_, 10 glucose, 5 KCl, 1.5 MgCl_2_, 2.5 CaCl_2_, 1.23 NaH_2_PO_4_; pH was titrated to 7.25 with continuously bubbling with 95% O_2_ with 95% CO_2_ for at least 20 minutes and osmolality was maintained at 305mOsm) for 30 minutes. After recovery, Cold-sensitive neurons in the lPBN (expressing fluorescence reporter protein) were recorded with a Multiclamp 700B amplifier (Axon Instruments) and Digidata-1550B (Axon Instruments), glass micropipettes electrode pulled from borosilicate glass capillaries (BF150-86-10, Sutter Instrument) with P-700 microprocessor-controlled puller (Sutter Instruments) containing K-gluconate based intracellular solution (containing (in mM) 123 K-gluconate, 18 KCl, 10 NaCl, 3 MgCl2, 2ATP-Na2, 0.3 GTP-Na, and 0.2 EGTA; adjusted to pH 7.3 with 1N KOH) in room-temperature aCSF. Resistance of glass micropipettes was kept at 3-5 MΩ. For the resting membrane potential recording, current clamp recording was performed, with holding current was held at 0 pA, 3 minute after membrane rupture for internal solution substitution. For the spontaneous action potential recording, recording was performed in current clamp mode, while holding current was held at 0 pA. Cells showing action potential firing during 60 seconds baseline recording periods were considered as spontaneous action potential firing neurons. For CNO perfusion experiment, after whole-cell recording configuration was successfully formed, we recorded 1 minutes of baseline period (during normal extracellular solution perfusion, 1.5mL/min of perfusion rate), followed by CNO (10μM) bath perfusion. After CNO perfusion, we recorded for additional 20 minutes as washout condition. Action potential firing frequency and average membrane potential was analyzed with Clampfit (v 10.7.0.3, Molecular Devices) in each time window. During the whole-cell patch clamp recording, cells with access resistance (R_a_) higher than 30 MΩ and/or membrane resistance lower than 250 MΩ were excluded from the analysis.

#### RNA sampling and sequencing analysis

Single cell suspension was sorted with FACSAria^TM^ III cell sorter (BD biosciences) with a 70nm nozzle. Forward scattering with Area (FSC-A) and side scattering with Area (SSC-A) parameters were used to isolate cells from debries, and singles were gated with Forward scattering with Area and width (FSC-A and FSC-W). After singlet gating, detection window of DAPI and ZsGreen fluorescent signal was set with the single cell suspension from wild-type mice, and fluorescent gates were compensation with single-and double-stained samples for each experiments. According to gating strategy, ZsGreen^+^, DAPI^-^ Cell populations (Zsg^+^ viable population) and ZsGreen^-^, DAPI^-^ Cell populations (Zsg^-^ viable population) were sorted into ice-cold aCSF containing 0.05% BSA. After sorting, collection tube was centrifuged at 660g for 5 minutes and supernatant was removed. Remaining cell pellets were dissolved with 2mL Trizol solution and stored at -80°C before RNA isolation. Total RNA was isolated using Trizol reagent (Invitrogen) or Maxwell® RSC miRNA from Kits (Promega™ Corporation). RNA quality was assessed by Agilent 4200 TapeStation System (Agilent Technologies), and RNA quantification was performed using Qubit (Thermo Fisher Scientific Inc.). Only RNA samples with RNA integrity (RIN) higher than 7.0 was included in further procedures. For control and test RNAs, the construction of library was performed using QuantSeq 3’ mRNA-Seq Library Prep Kit FWD (Lexogen) according to the manufacturer’s instructions. In brief, each total RNA were prepared and an oligo-dT primer containing an Illumina-compatible sequence at its 5’ end was hybridized to the RNA and reverse transcription was performed. After degradation of the RNA template, second strand synthesis was initiated by a random primer containing an Illumina compatible linker sequence at its 5’ end. The double-stranded library was purified by using magnetic beads to remove all reaction components. The library was amplified to add the complete adapter sequences required for cluster generation. The finished library is purified from PCR components. High-throughput sequencing was performed as single-end 75bp sequencing using NextSeq 500 / 550 (Illumina Inc.). A quality control of raw sequencing data was performed using FastQC. Sequenced reads were trimmed for adapter sequences and low-quality filtered using bbduk. Then the clean reads were mapped to the reference genome using STAR (*77*). The quantification of reads was processed using HTSeq-count (*78*). The Read Counts were processed based on TMM+CPM normalization method using Python “conorm” package. Data mining and graphic visualization were performed using ExDEGA (Ebiogen Inc., Korea). Genes were considered significantly enriched if they exhibited a fold change > 2 and have a p-value < 0.05 in Zsg-positive cells compared to Zsg-negative cells, while those with a fold change of < 0.5 were regarded as significantly depleted.

#### In situ hybridization

RNAscope multiplex v2 kit (Advanced Cell Diagnostics (ACD)) was used to perform fluorescence in situ hybridization according to the manufacturer’s protocols. For tissue preparation, mice were undergone trans-cardiac perfusion with ice-cold Phosphate-buffered saline (PBS) treated with Diethyl Pyrocarbonate (DEPC) under anesthesia with sodium pentobarbital (55mg/kg). The harvested brain tissues were embedded in OCT compound, rapidly frozen and cryosectioned by 12 µm per section and stored at -80°C before in situ hybridization. On the day of the fluorescence in situ hybridization experiment, slides were directly fixed in 4% paraformaldehyde (PFA) for 1 hour followed by wash with DEPC-treated PBS and dehydration in 50%, 75% and 100% ethanol at room temperature. Tissue samples were then processed followed by RNAscope Multiplex fluorescent reagent kit v2 protocol (ACD, Document Number UM323100). Slides were imaged with Axioscan Slide scanner and/or LSM 980 confocal microscopy (40x) then analyzed with QuPath v 0.3.2. Commercial probes used in fluorescence in situ hybridization is listed below.

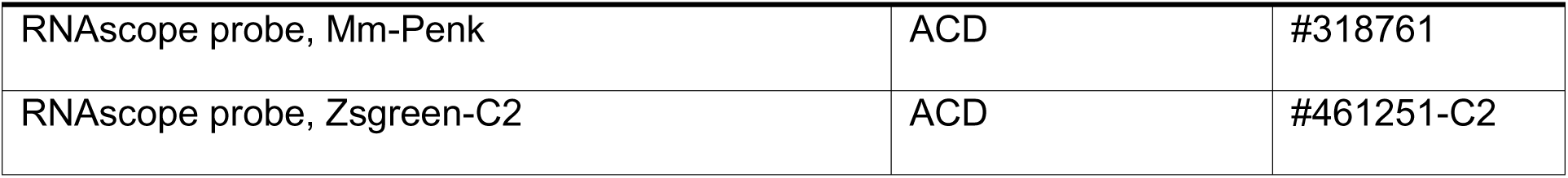

|  |  |  |
| --- | --- | --- |
| RNAscope probe, Mm-Penk | ACD | #318761 |
| RNAscope probe, Zsgreen-C2 | ACD | #461251-C2 |

#### Statistical analysis

Comparison between two groups was analyzed by Student’s *t*-test (paired or unpaired), and comparison between multiple groups was analyzed by ANOVA and Bonferroni’s *post hoc* tests or Dunnett post-tests using Prism software (version 8.4.2, GraphPad). Detailed statistical methods and descriptive statistics were provided with supplementary tables. Statistical significance was set at *P* < 0.05, and all data are presented as mean ± SEM.

