## Supplementary Figures 1-8 for "Brainstem enkephalinergic neural circuit underlying cold-induced pain relief in mice"

**Figure S1: Effect of cold in mechanical allodynia and heat hyperalgesia in CFA-induced inflammatory pain**

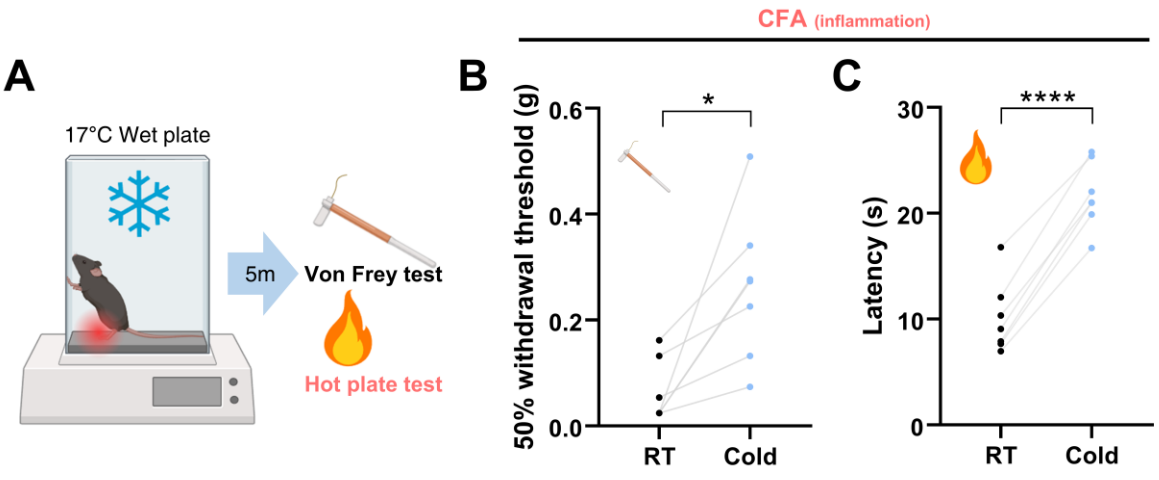


(**A**) Experimental scheme for the behavior test presented in Figure S1B-C.

(**B**) Mechanical sensitivity measure with 50% withdrawal thresholds in CFA-induced inflammatory pain model mice in room temperature and Cold treatment.

(**C**) Thermal sensitivity to noxious hot measured with response latency to hot plate in CFA-induced inflammatory pain model mice in room temperature and Cold treatment.

Data are presented as mean ± SEMs. $*p < 0.05$, $**p < 0.01$, $***p < 0.001$ and $****p < 0.0001$.

Number of mice and statistical tests are listed in **Table S1**.

**Figure S2: Effect of icilin in membrane depolarization-induced calcium response in peripheral sensory neurons**

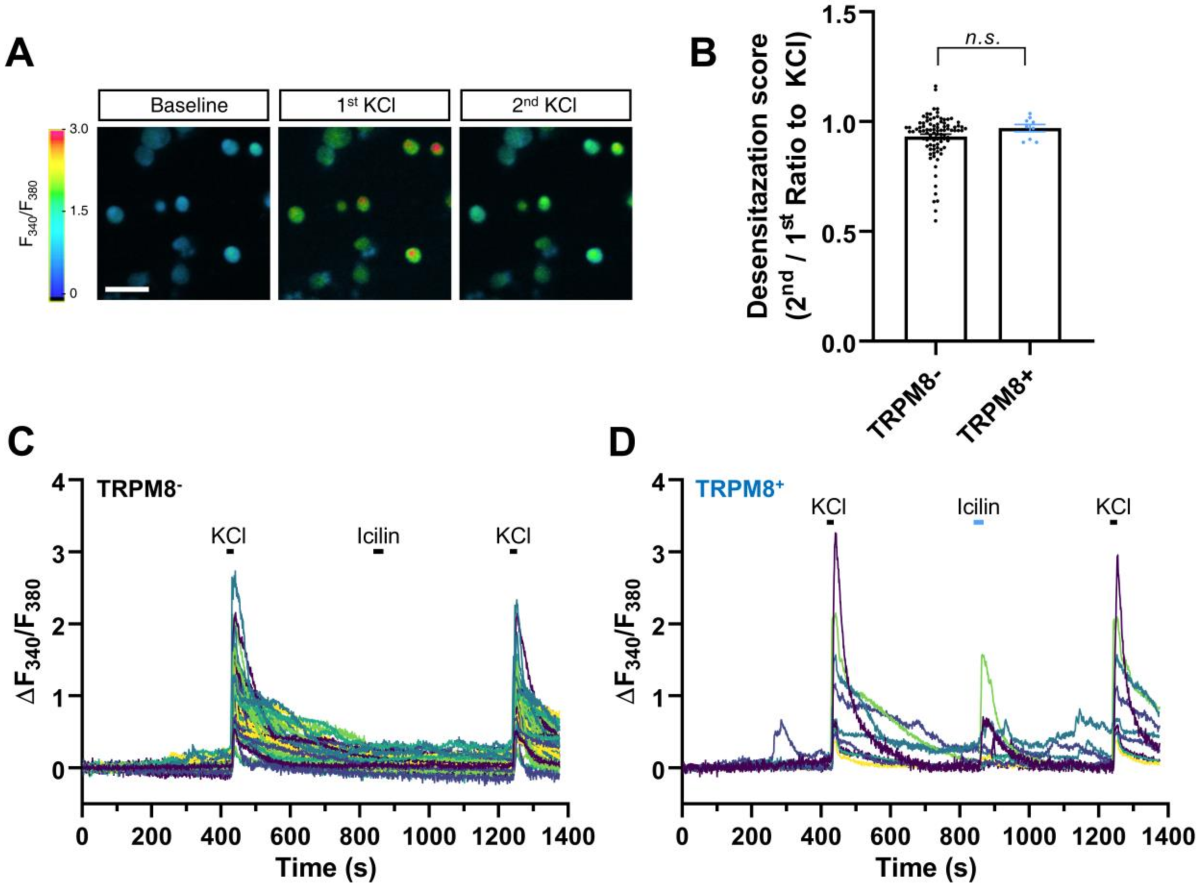


(**A**) Representative ratiometric image of Fura-2 calcium dye in primary cultured DRG neurons.

(**B**) Quantification of desensitization score to KCl response in TRPM8$^+$ and TRPM8$^-$ DRG neurons.

(**C**) Representative calcium traces from TRPM8$^-$ DRG neurons.

(**D**) Representative calcium traces from TRPM8$^+$ DRG neurons.

Data are presented as mean ± SEMs. $*p < 0.05$, $**p < 0.01$, $***p < 0.001$ and $****p < 0.0001$. Number of mice and statistical tests are listed in **Table S1**.

**Figure S3: Lack of CGRP expression in lPBN$^{cold}$ neurons**

**A**

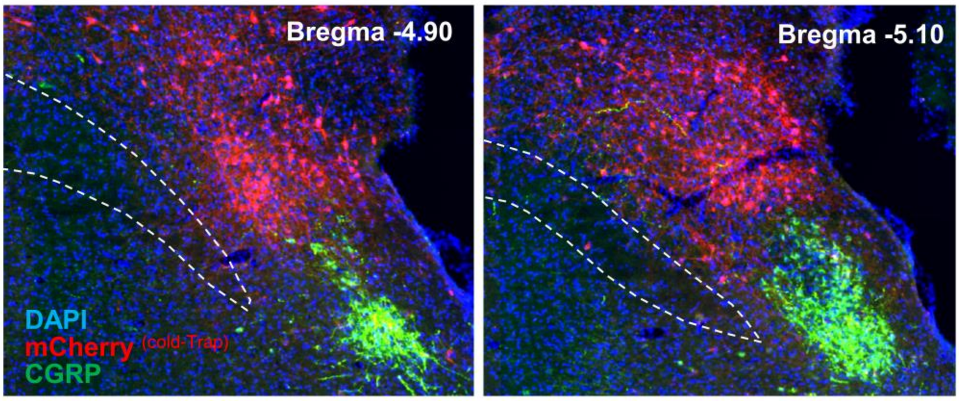


(**A**) Distribution of mCherry-positive (Cold-TRAP) neurons and anti-CGRP immunostaining in lateral parabrachial nucleus. Green; CGRP. Red; mCherry (cold-TRAP).

**Figure S4: Double labeling experiments in lateral parabrachial nucleus for different stimuli**

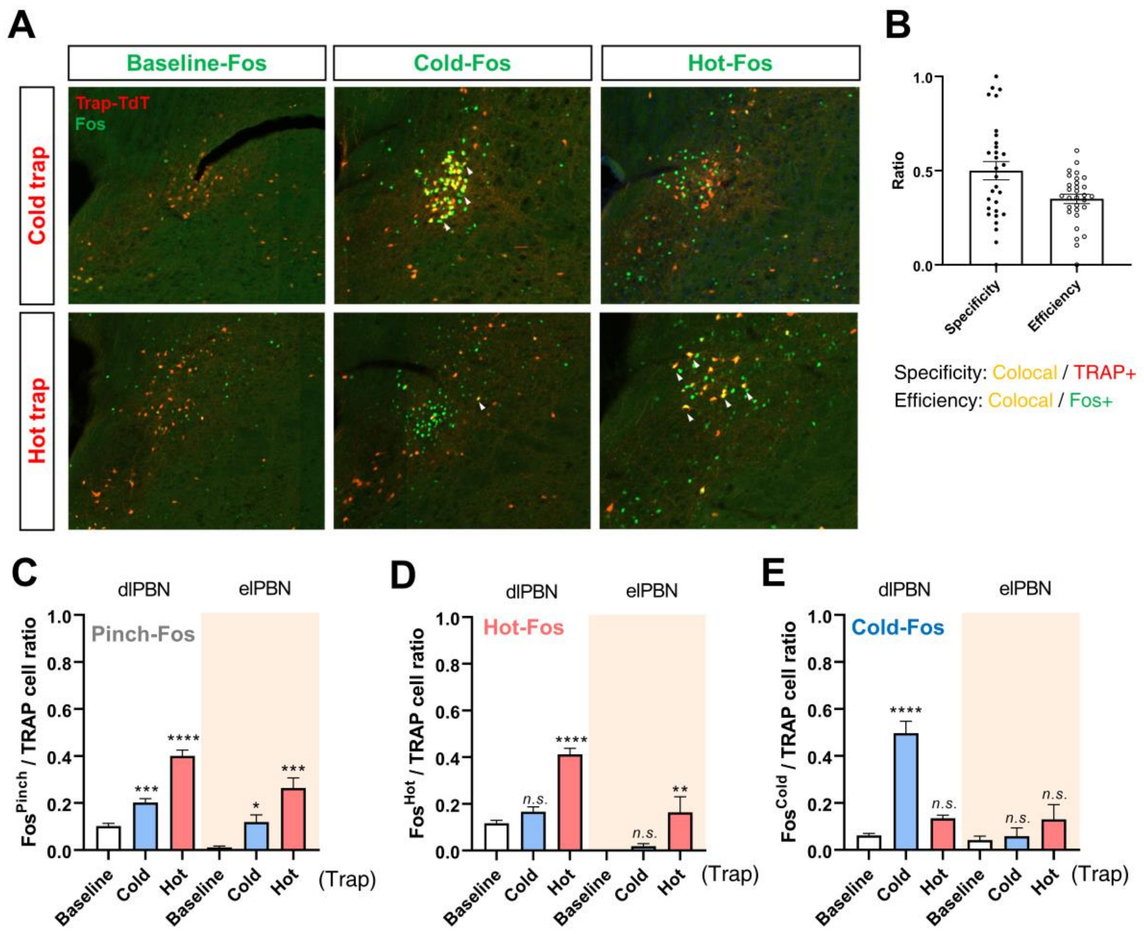


(**A**) Representative immunohistochemical images show neurons activated and labeled with TdTomato (TRAP signal) and c-Fos in the lateral parabrachial nucleus, with different stimuli

(**B**) Quantification of specificity and efficiency of TRAP methods.

(**C**) Ratio of pinch-Fos-positive cells in baseline-,cold-,hot-TRAP lateral parabrachial neurons.

(**D**) Ratio of hot-Fos-positive cells in baseline-,cold-,hot-TRAP lateral parabrachial neurons.

(**E**) Ratio of cold-Fos-positive cells in baseline-,cold-,hot-TRAP lateral parabrachial neurons.

Data are presented as mean ± SEMs. $*p < 0.05$, $**p < 0.01$, $***p < 0.001$ and $****p < 0.0001$. Number of mice and statistical tests are listed in **Table S1**.

**Figure S5: Electrophysiological validation of DREADD and characterization of lPBN$^{cold}$ neurons**

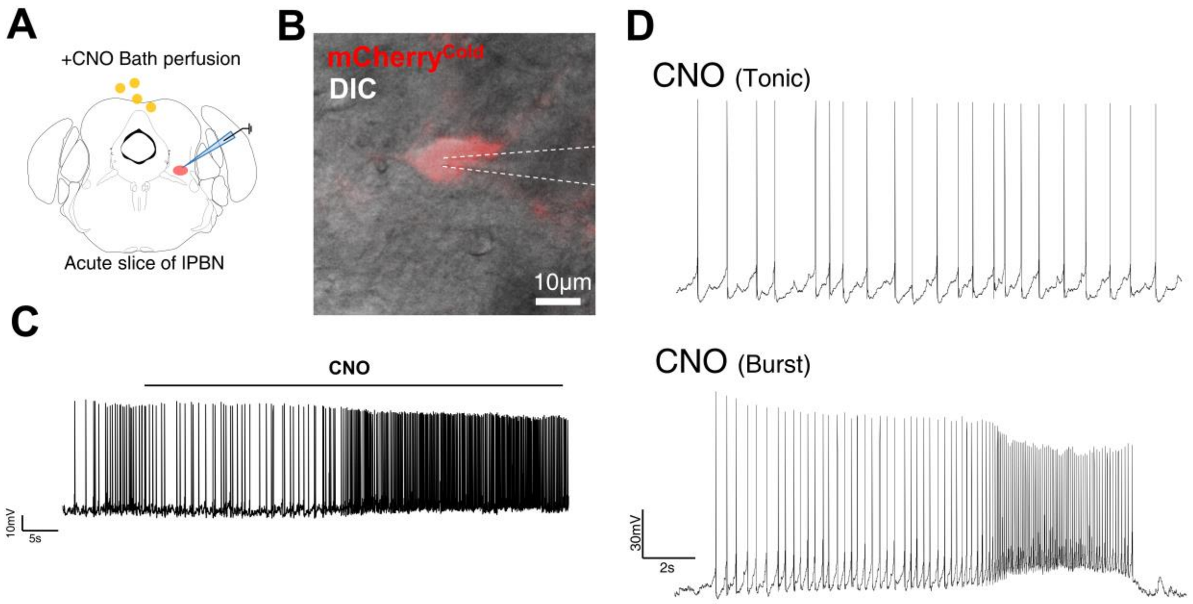


(**A**) Experimental scheme for the eletrophysiological recording presented in Figure S5.

(**B**) Representative image of mCherry-positive lPBN$^{cold}$ neurons. Red; mCherry.

(**C**) Representative membrane potential traces of lPBN$^{cold}$ neurons during CNO treatment (10μM).

(**D**) Representative membrane potential traces of lPBN$^{cold}$ neurons during CNO treatment.

Data are presented as mean ± SEMs. Number of mice and statistical tests are listed in **Table S1**.

**Figure S6: Activation of lPBN$^{cold}$ neurons increase spontaneous cold avoidance**

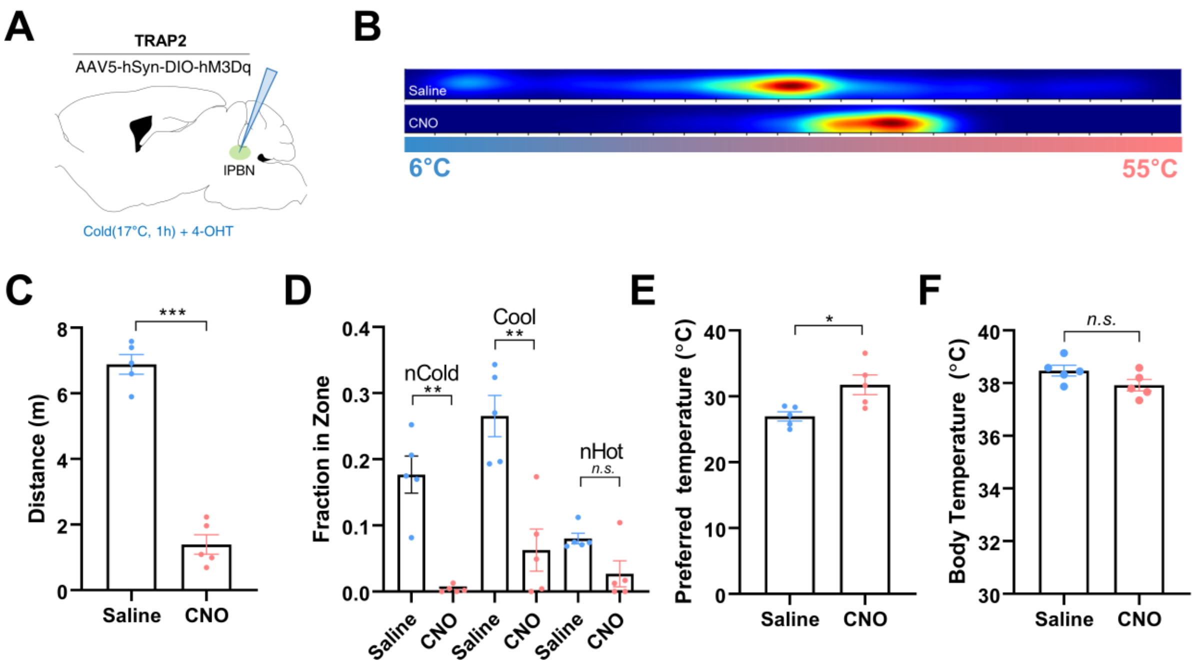


(**A**) Experimental scheme for the behavior test presented in Figure S6B-F.

(**B**) Heatmap plot for thermal gradient test in saline-treated (top) and CNO-treated (bottom) mice.

(**C**) Total distance moved in thermal gradient test.

(**D**) Fraction of time spent in each Zone of thermal gradient plate.

(**E**) Preferred temperature of mice in thermal gradient test.

(**F**) Core body temperature of mice, measure with rectal probe.

Data are presented as mean ± SEMs. $*p < 0.05$, $**p < 0.01$, $***p < 0.001$ and $****p < 0.0001$.

Number of mice and statistical tests are listed in **Table S1**.

**Figure S7: Activation of vlPAG-projecting lPBN neurons to cold**

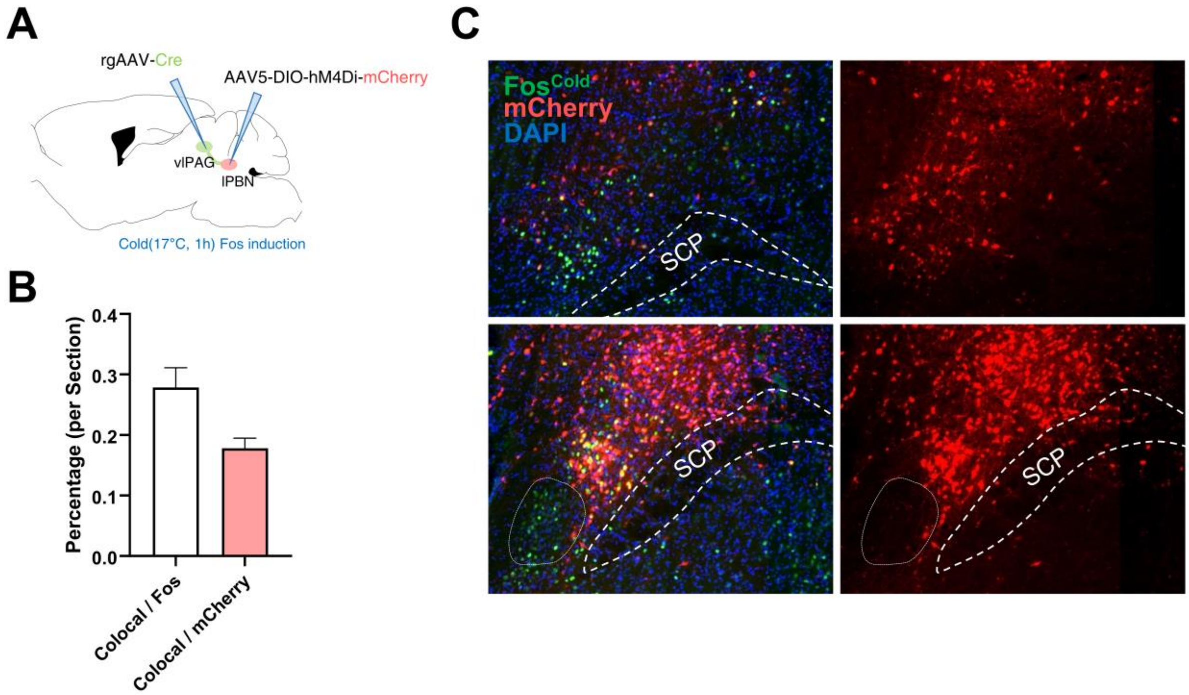


(**A**) Experimental scheme for the AAV-mediated tracing of vlPAG-projecting lPBN neurons.

(**B**) Quantification of cellular responsiveness.

(**C**) Representative immunohistochemical images show vlPAG-projecting lPBN neurons labeled with TdTomato and cold-induced c-Fos in the lateral parabrachial nucleus.

Data are presented as mean ± SEMs. Number of mice and statistical tests are listed in **Table S1**.

**Figure S8: Ablation of IPBN$^{cold}$ neurons does not affect thermal preference, basal body temperature and baseline nociception**

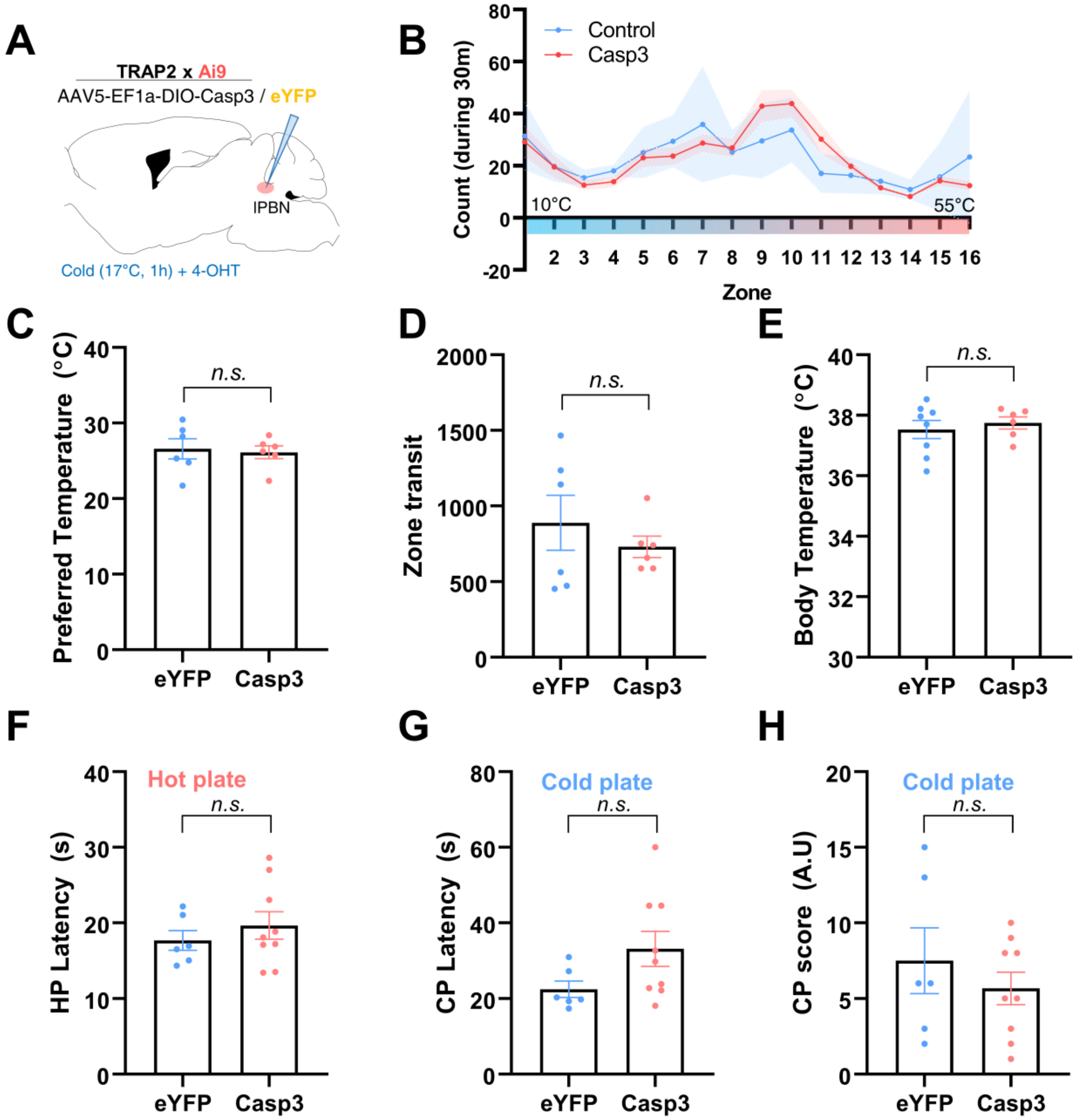


(**A**) Experimental scheme for the AAV-mediated selective ablation of IPBN$^{cold}$ neurons.

(**B**) Quantification of thermal gradient test in control mice (Blue) and AAV-mediated ablation (Red) mice.

(**C**) Preferred temperature of mice in thermal gradient test.

(**D**) Total zone transit in thermal gradient test.

(**E**) Core body temperature of mice, measure with rectal probe.

(**F**) Thermal sensitivity to noxious hot measured with response latency to hot plate in lPBN$^{cold}$ neuron activation experiment.

(**G**) Thermal sensitivity to noxious cold measured with response latency to cold plate in lPBN$^{cold}$ neuron activation experiment.

(**H**) Thermal sensitivity to noxious cold measured with nocifensive score to cold plate in lPBN$^{cold}$ neuron activation experiment.

Data are presented as mean ± SEMs. $*p < 0.05$, $**p < 0.01$, $***p < 0.001$ and $****p < 0.0001$. Number of mice and statistical tests are listed in **Table S1**.

### Supplementary Table 1: Number of mice/neurons and statistical tests in main figures and Supplementary Figures

| Fig. | Number of experimental objects | Statistical test | Descriptive statistics ( Mean ± SEM) |
|---|---|---|---|
| 1B | Vehicle and icilin, *n* = 15 mice each | Paired t-test, P = 0.0297 | 0.5880 ± 0.0574 for vehicle, 0.9460 ± 0.1310 for icilin |
| 1C | Vehicle and icilin, *n* = 8 mice each | Paired t-test, P = 0.0027 | 0.1162 ± 0.031 for vehicle, 0.778 ± 0.173 for icilin |
| 1D | Vehicle and icilin, *n* = 5 mice each | Paired t-test, P = 0.0113 | 0.0271 ± 0.0052 for vehicle, 0.287 ± 0.0573 for icilin |
| 1E, bottom | Control TRAP, *n* = 3 mice; Cold TRAP, *n* = 7 mice | Paired t-test, P = 0.0033 for ACC, P = 0.485 for aIC, P < 0.0001 for pIC, P < 0.0001 for S1, P = 0.477 for PVT, P = 0.477 for DMH, P = 0.516 for VMH, P = 0.0166 for vlPAG | 1.38 ± 0.179 for ACC, 1.03 ± 0.374 for aIC, 2.224 ± 0.654 for pIC, 2.096 ± 0.478 for S1, 0.909 ± 0.279 for PVT, 1.174 ± 0.338 for DMH, 0.823 ± 0.372 for VHM, 1.742 ± 0.421 for vlPAG (fold change) |
| 1G, left | Control TRAP, *n* = 3 mice; Cold TRAP, *n* = 7 mice | Paired t-test, P < 0.0001 for -4.9mm, P = 0.805 for -5.1mm, P = 0.770 for -5.3mm | -4.9mm: 3.50 ± 0.69 for Control, 12.74 ± 1.42 for Cold / -5.10mm; 2.30 ± 0.60 for control, 2.10 ± 0.403 for Cold / -5.30mm; 0.53 ± 0.355 for control, 0.66 ± 0.245 for cold |
| 1G, right | Control TRAP, *n* = 3 mice; Cold TRAP, *n* = 7 mice | Paired t-test, P = 0.838 for -4.9mm, P = 0.048 for -5.1mm, P = 0.282 for -5.3mm | -4.9mm: 15.67 ± 2.68 for Control, 14.91 ± 2.15 for Cold / -5.10mm; 17.40 ± 1.93 for control, 23.66 ± 1.62 for Cold / -5.30mm; 4.88 ± 1.37 for control, 6.83 ± 1.07 for cold |
| 2E, right | *n* = 16 cells each | Unpaired t-test, P = 0.0014 | 0.379 ± 0.113 for $LPBN^{HP}$ neurons, -0.173 ± 0.107 for $LPBN^{cold}$ neurons |
| 2F, right | *n* = 16 cells each | Unpaired t-test, P < 0.0001 | 0.779 ± 0.151 for $LPBN^{HP}$ neurons, -0.360 ± 0.192 for $LPBN^{cold}$ neurons |
| 2G, right | *n* = 16 cells each | Unpaired t-test, P = 0.0001 | -0.603 ± 0.126 for $LPBN^{HP}$ neurons, 0.2991 ± 0.165 for $LPBN^{cold}$ neurons |
| 3C | *n* = 7 cells from 4 mice | One-Way ANOVA with Dunnett's multiple comparisons test, P = 0.0400 for Baseline vs CNO, P = 0.0665 for Bath vs washout. | 0.531 ± 0.052 for Saline-treated, 1.494 ± 0.191 for CNO (0.1mg/kg) treated, 2.129 ± 0.221 for CNO (1mg/kg) treated |

| | | | |
|---|---|---|---|
| 3E | *n* = 10 mice for Saline-treated and CNO (1mg/kg) treated, n = 5 mice for CNO (0.1mg/kg) treated | One-Way ANOVA with Dunnett's multiple comparisons test, P < 0.0001 for Treatment; P = 0.164 for Saline vs CNO (0.1mg/kg), P < 0.0001 for Saline vs CNO (1mg/kg) | 0.535 ± 0.303 for baseline, 2.355 ± 0.850 for CNO, 0.996 ± 0.423 fop washout |
| 3F | *n* = 10 mice for Saline-treated and CNO (1mg/kg) treated, n = 5 mice for CNO (0.1mg/kg) treated | One-Way ANOVA with Dunnett's multiple comparisons test, P < 0.0001 for Treatment; P = 0.164 for Saline vs CNO (0.1mg/kg), P < 0.0001 for Saline vs CNO (1mg/kg) | 12.52 ± 1.03 for Saline-treated, 16.21 ± 1.29 for CNO (0.1mg/kg) treated, 19.60 ± 0.245 for CNO (1mg/kg) treated |
| 3G | *n* = 15 mice for Saline-treated and CNO (1mg/kg) treated, n = 5 mice for CNO (0.1mg/kg) treated | One-Way ANOVA with Dunnett's multiple comparisons test; P = 0.532 for Saline vs CNO (0.1mg/kg), P = 0.048 for Saline vs CNO (1mg/kg) | 27.65 ± 2.87 for Saline-treated, 22.56 ± 6.87 for CNO (0.1mg/kg) treated, 16.72 ± 3.41 for CNO (1mg/kg) treated |
| 3H | *n* = 15 mice for Saline-treated and CNO (1mg/kg) treated, n = 5 mice for CNO (0.1mg/kg) treated | One-Way ANOVA with Dunnett's multiple comparisons test, P = 0.0046 for Treatment; P = 0.0404 for Saline vs CNO (0.1mg/kg), P = 0.0014 for Saline vs CNO (1mg/kg) | 6.87 ± 0.96 for Saline-treated, 17.80 ± 4.50 for CNO (0.1mg/kg) treated, 20.13 ± 3.31 for CNO (1mg/kg) treated |
| 4D | *n* = 4 mice each | Paired t-test, P = 0.0393 | 0.480 ± 0.0091 for vehicle, 1.134 ± 0.106 for CNO |
| 4E | *n* = 4 mice each | Paired t-test, P = 0.0184 | 18.47 ± 1.94 for vehicle, 26.15 ± 1.765 for CNO |
| 4F | *n* = 4 mice each | Paired t-test, P = 0.0242 | 13.05 ± 2.77 for vehicle, 2.790 ± 0.835 for CNO |
| 4G | *n* = 4 mice each | Paired t-test, P = 0.0017 | 14.50 ± 3.23 for vehicle, 37.50 ± 1.56 for CNO |
| 4H, left | *n* = 4 mice each | Paired t-test, P = 0.486 | 11.10 ± 1.977 for vehicle, 14.31 ± 2.885 for CNO |
| 4H, right | *n* = 4 mice each | Paired t-test, P = 0.0017 | 17.93 ± 2.13 for vehicle, 24.28 ± 9.09 for CNO |
| 4I | *n* = 8 mice each | One-Way ANOVA with Dunnett's multiple comparisons test; P = 0.0486 for Saline vs CNO, P = 0.6858 for Saline vs CNO+Naloxone | 0.468 ± 0.056 for Saline, 0.961 ± 0.145 for CNO, 0.628 ± 0.172 for CNO+Nal |
| 4J | *n* = 5 mice each | One-Way ANOVA with Tukey's multiple comparisons test; P = 0.0095 for Baseline vs CFA-saline, P = 0.0.42 for CFA-saline vs CFA-CNO | 0.569 ± 0.0087 for Baseline, 0.0497 ± 0.028 for CFA-Saline, 1.611 ± 0.370 for CFA-CNO |

| | | | |
|---|---|---|---|
| 5C | *n* = 2 mice for Control, *n* = 3 mice for Casp3; 18 sections from Control; 30 sections from Casp3 | Unpaired t-test, P < 0.0001 | 30.44 ± 3.29 for Control, 9.90 ± 1.87 for Casp3 |
| 5D | *n* = 6 mice for control, *n* = 9 mice for Casp3 | Unpaired t-test, P = 0.917 | 0.630 ± 0.0237 for Control, 0.682 ± 0.078 for Casp3 |
| 5E | *n* = 5 mice each | Paired t-test, P = 0.5600 | 0.782 ± 0.056 for Baseline, 0.724 ± 0.074 for Vehicle |
| 5F | *n* = 15 mice each for control, *n* = 8 mice each for Casp3 | Paired t-test, P = 0.0297 for Control group, P = 0.7673 for Casp3 group | Control: 0.588 ± 0.057 for vehicle, 0.946 ± 0.131 for icilin; Casp3: 0.710 ± 0.051 for vehicle, 0.688 ± 0.067 for icilin |
| 5G, control | *n* = 8 mice each | 2 Way ANOVA with Sidak's multiple comparison test, Baseline; P = 0.7304, CFA; P = 0.0022, CFA+icilin; P = 0.9768 | Contralatearal: 0.680 ± 0.134 for Baseline, 0.775 ± 0.133 for CFA, 0.844 ± 0.132 for CFA+icilin; Ipsilateral: 0.510 ± 0.076 for Basline, 0.116 ± 0.0288 for CFA, 0.777 ± 0.161 for CFA+icilin |
| 5G, Casp3 | *n* = 8 mice each | 2 Way ANOVA with Sidak's multiple comparison test, Baseline; P = 0.6995, CFA; P = 0.6485, CFA+icilin; P = 0.0100 | Contralatearal: 0.805 ± 0.105 for Baseline, 0.956 ± 0.226 for CFA, 0.876 ± 0.088 for CFA+icilin; Ipsilateral: 0.614 ± 0.077 for Basline, 0.307 ± 0.105 for CFA, 0.274 ± 0.107 for CFA+icilin |
| 5H | *n* = 8 mice for control, *n* = 8 mice for Casp3 | Unpaired t-test, P = 0.0341 | 0.6613 ± 0.146 for control, 0.020 ± 0.242 for Casp3 |
| 5J | *n* = 8 mice each | Paired t-test; P = 0.0411 for saline, P = 0.4041 for CNO | Saline: 0.412 ± 0.0638 for vehicle, 0.750 ± 0.122 for icilin; CNO: 0.425 ± 0.0383 for vehicle, 0.4923 ± 0.0778 for icilin |
| 5c | *n* = 4 expreimental batches for WT, *n* = 5 batches for TRAP-ZsGreen (each with 4-5 mice) | Unpaired t-test, P = 0.0082 | 0.2750 ± 0.103 for WT, 5.880 ± 1.354 for TRAP |
| 6B | *n* = 5 batches | N.A. | 79.34 ± 2.665 (%) |
| 6I | *n* = 2 mice for Saline, *n* = 3 mice for CNO | Unpaired t-test, P = 0.0054 | 1.00 ± 0.034 for Control, 1.11 ± 0.0157 for cold |
| 6J | *n* = 3 mice each | Unpaired t-test, P = 0.0414 | 1.00 ± 0.017 for Control, 1.054 ± 0.0175 for cold |
| 6L | *n* = 6 mice each | Paired t-test, P = 0.0481 for Saline-treated, P = 0.0924 for anti-met-enk treated | 0.0156 ± 0.0232 for Saline-vehicle, 0.1690 ± 0.0287 for saline-icilin, 0.0456 ± 0.0176 for anti-enk-vehicle, 0.1068 ± 0.0334 for anto-enk-icilin |

| | | | |
|---|---|---|---|
| 7A, right | Control TRAP, *n* = 3 mice; Cold TRAP, *n* = 7 mice | Unpaired t-test, P < 0.0001 | 25.82 ± 2.085 for Control, 49.42 ± 3.07 for Cold |
| 7B, right | Control TRAP, *n* = 3 mice; Cold TRAP, *n* = 7 mice | Unpaired t-test, P = 0.0104 | 8.40 ± 1.22 for Control, 12.78 ± 0.96 for Cold |
| 7E | *n* = 2 mice for Saline, *n* = 3 mice for CNO | Unpaired t-test, P < 0.0001 | 19.91 ± 0.665 for Saline, 11.83 ± 0.653 for CNO |
| 7G | *n* = 2 mice for Saline, *n* = 3 mice for CNO | Unpaired t-test, P < 0.0001 | 43.93 ± 1.06 for Saline, 32.59 ± 1.43 for CNO |
| 8C | *n* = 5 mice each | Paired t-test, P = 0.0348 | 0.474 ± 0.063 for saline, 1.741 ± 0.387 for CNO |
| 8D | *n* = 5 mice each | Paired t-test, P < 0.0001 | 16.66 ± 0.837 for saline, 29.73 ± 0.271 for CNO |
| 8E | *n* = 5 mice each | Paired t-test, P = 0.2545 | 6.20 ± 2.15 for saline, 3.80 ± 1.16 for CNO |
| 8H | *n* = 2 mice for Saline, *n* = 3 mice for CNO | Unpaired t-test, P < 0.0001 | 13.64 ± 0.569 for Saline, 6.05 ± 0.207 for CNO |
| 8J | *n* = 2 mice for Saline, *n* = 3 mice for CNO | Unpaired t-test, P < 0.0001 | 35.47 ± 1.088 for Saline, 22.82 ± 0.5258 for CNO |
| **Supple. Fig** | **Number of experimental objects** | **Statistical test** | **Descriptive statistics ( Mean ± SEM)** |
| 1B | *n* = 7 mice each | Paired t-test, P = 0.0130 | 0.063 ± 0.0221 for RT, 0.261 ± 0.0537 for Cold |
| 1C | *n* = 7 mice each | Paired t-test, P < 0.0001 | 10.11 ± 1.295 for RT, 21.68 ± 1.190 for Cold |
| 2B | 92 $TRPM8^-$ Cells, 9 $TRPM8^+$ Cells | Unpaired t-test, P = 0.2382 | 0.9324 ± 0.010 for TRPM8- Cells, 0.9702 ± 0.0164 for TRPM8+ Cells |
| 4B | 30 sections each, from 5 mice | N.A. | specificity of 0.500 ± 0.048, efficiency of 0.350 ± 0.135 |
| 4C | *n* = 5 mice each; 39 sections from baseline TRAP; 38 sections from Cold TRAP; 36 sections from Hot trap | One-Way ANOVA with Dunnett's multiple comparisons test,P < 0.0001 for Baseline vs | 0.102 ± 0.012 for Baseline, 0.400 ± 0.0253 for Cold, 0.202 ± 0.017 for Hot (dlPBN); 0.011 ± 0.0061 for Baseline, 0.264 ± |

| | | | |
|---|---|---|---|
| | | Cold, P = 0.0004 for Baseline vs Hot (dlPBN); P < 0.0001 for Baseline vs Cold, P = 0.0180 for Baseline vs Hot (elPBN) | 0.043 for Cold, 0.120 ± 0.030 for Hot (elPBN) |
| 4D | *n* = 4 mice each; 25 sections from baseline TRAP; 28 sections from Cold TRAP; 23 sections from Hot trap | One-Way ANOVA with Dunnett's multiple comparisons test, P = 0.1542 for Baseline vs Cold, P < 0.0001 for Baseline vs Hot (dlPBN); P = 0.8834 for Baseline vs Cold, P = 0.0022 for Baseline vs Hot (elPBN) | 0.1172 ± 0.012 for Baseline, 0.1659 ± 0.021 for Cold, 0.412 ± 0.026 for Hot (dlPBN); 0 for Baseline, 0.0183 ± 0.011 for Cold, 0.1633 ± 0.067 for Hot (elPBN) |
| 4E | *n* = 4 mice each; 28 sections from baseline TRAP; 30 sections from Cold TRAP; 22 sections from Hot trap | One-Way ANOVA with Dunnett's multiple comparisons test,P < 0.0001 for Baseline vs Cold, P = 0.2217 for Baseline vs Hot (dlPBN); P = 0.9244 for Baseline vs Cold, P = 0.2245 for Baseline vs Hot (elPBN) | 0.042 ± 0.017 for Baseline, 0.058 ± 0.035 for Cold, 0.130 ± 0.063 for Hot (dlPBN); 0.062 ± 0.0084 for Baseline, 0.497 ± 0.051 for Cold, 0.135 ± 0.013 for Hot (elPBN) |
| 6C | *n* = 5 mice each | Paired t-test, P = 0.0004 | 6.880 ± 0.300 for Saline, 1.393 ± 0.298 for CNO |
| 6D | *n* = 5 mice each | Paired t-test, P = 0.0028 (noxious cold Zone); P = 0.0099 (innocuous cold Zone); P = 0.0803 (noxious hot Zone) | Saline; 0.177 ± 0.028 for Noxious cold, 0.2654 ± 0.0312 for innocuous cold, 0.0802 ± 0.0081 for noxious hot; CNO; 0.0038 ± 0.0024 for Noxious cold, 0.0627 ± 0.032 for innocuous cold, 0.0267 ± 0.0197 for noxious hot |
| 6E | *n* = 5 mice each | Paired t-test, P = 0.0188 | 26.96 ± 0.695 for Saline, 31.75 ± 1.49 for CNO |
| 6F | *n* = 5 mice each | Paired t-test, P = 0.0651 | 38.47 ± 0.204 for Saline, 37.92 ± 0.216 for CNO |
| 7B | From 3 mice, 10 sections each | N.A. | 0.273 ± 0.033 for colocal / Fos, 0.178 ± 0.0167 for Colocal / mCherry |
| 8C | *n* = 6 mice each | Unpaired t-test, P = 0.7842 | 26.58 ± 1.318 for Control, 26.13 ± 0.840 for Casp3 |
| 8D | *n* = 6 mice each | Unpaired t-test, P = 0.4988 | 888.5 ± 181.5 for Control, 730 ± 70.67 for Casp3 |
| 8E | *n* = 6 mice each | Unpaired t-test, P = 0.5910 | 37.53 ± 0.302 for Control, 37.74 ± 0.201 for Casp3 |

| | | | |
|---|---|---|---|
| 8F | *n* = 6 mice each | Unpaired t-test, P = 0.4422 | 17.69 ± 1.31 for Control, 19.66 ± 1.818 for Casp3 |
| 8G | *n* = 6 mice each | Unpaired t-test, P = 0.0976 | 22.46 ± 2.182 for Control, 33.15 ± 4.618 for Casp3 |
| 8H | *n* = 6 mice each | Unpaired t-test, P = 0.4187 | 7.50 ± 2.172 for Control, 5.667 ± 1.08 for Casp3 |
